# SUMOylation controls the rapid transcriptional reprogramming induced by anthracyclines in Acute Myeloid Leukemias

**DOI:** 10.1101/2022.04.19.488613

**Authors:** Mathias Boulanger, Chamseddine Kifagi, Marko Ristic, Ludovic Gabellier, Denis Tempé, Jon-Otti Sigurdsson, Tony Kaoma, Charlotte Andrieu-Soler, Thierry Forné, Eric Soler, Yosr Hicheri, Elise Gueret, Laurent Vallar, Jesper V Olsen, Guillaume Cartron, Marc Piechaczyk, Guillaume Bossis

## Abstract

Genotoxicants have been used for decades as front-line therapies against cancer on the basis of their DNA-damaging actions. However, some of their non-DNA-damaging effects are also instrumental for killing dividing cells. We report here that the anthracycline Daunorubicin (DNR), one of the main drugs used to treat Acute Myeloid Leukemia (AML), induces broad transcriptional changes in AML cells before cell death induction. The regulated genes are particularly enriched in genes controlling cell proliferation and death, as well as inflammation and immunity. These transcriptional changes are preceded by DNR-dependent *de*SUMOylation of chromatin proteins, which limits both the positive and negative effects of DNR on transcription. Quantitative proteomics shows that proteins that are *de*SUMOylated in response to DNR are mostly transcription factors, transcriptional co-regulators and chromatin organizers. Among them, the CCCTC-binding factor CTCF is highly enriched at SUMO-binding sites found in *cis*-regulatory regions. This is notably the case at the promoter of the DNR-induced *NFKB2* gene. Its induction is preceded by a SUMO-dependent reconfiguration of chromatin loops engaging its CTCF- and SUMO-bound promoter with distal *cis*-regulatory regions. Altogether, our work suggests that one of the earliest effects of DNR in AML cells is a SUMO-dependent transcriptional reprogramming.

## Introduction

Acute Myeloid Leukemias (AML) are severe hematological malignancies, which arise through the acquisition of oncogenic mutations by hematopoietic stem- or progenitor cells from the myeloid lineage. Although AML constitutes a highly heterogenous group of diseases, most of them are treated similarly with the combination of one anthracycline, such as Daunorubicin (DNR), and the nucleoside analogue Cytarabine (Ara-C) (1–3). Most patients respond to this remission induction treatment (most often just called “induction treatment”). However, a large fraction of them relapse and become refractory to the drugs, which contributes to the dismal prognosis of this disease (2, 3). It is therefore critical to better understand the mode(s) of action of these drugs to find ways to overcome chemoresistance.

The DNA-damaging properties of both Ara-C and DNR are essential for therapeutical efficacy and have been characterized extensively (4, 5). However, these drugs also display many other cellular effects that can both favor or counteract their ability to induce cell death, even though these have little been studied so far. For example, anthracyclines can induce fast production of reactive oxygen species (ROS) that contribute to apoptosis induction by activating various signaling pathways (6). On the other hand, Ara-C and DNR also activate, at the same time, many pro-survival pathways that mitigate their pro-apoptotic actions. This is notable for the PI3K/AKT- (7), MAPK- (8) and NF-κB (9, 10) pathways, as their inhibitions potentiate genotoxics-induced cell death in cancer cells. Finally, both anthracyclines and Ara-C have long been known to alter transcriptional programs on the mid/long term (day-range) when used at sublethal doses (11, 12). However, how Ara-C and DNR contribute to gene expression changes, in particular at early times after the start of a treatment, has been poorly investigated.

We have formerly shown that one early consequence of DNR and Ara-C treatments is ROS-dependent *de*SUMOylation of cellular proteins in chemosensitive AMLs, which participates in induction of apoptosis (13). SUMOylation consists of reversible, covalent modification of cell proteins by the ubiquitin-related peptidic post-translational modifiers SUMO-1 to -3. SUMO-1 is 50% identical to SUMO-2 and -3, which are 95% identical and frequently referred to as SUMO-2/3 as their individual functions can often not be distinguished. The three SUMOs are conjugated by a conserved enzymatic cascade comprising one SUMO-activating enzyme (SAE1/SAE2 dimer; also called SUMO E1), one SUMO-conjugating enzyme (Ubc9; also called SUMO E2) and several SUMO E3s that facilitate SUMO transfer from the E2 onto its protein targets. SUMOylation is highly dynamic thanks to various isopeptidases (also called *de*SUMOylases) that remove SUMO from its substrates (14). Thousands of SUMOylated proteins involved in many cellular processes have now been identified (15). However, one of the main biological processes associated with SUMOylation is the control of gene expression. Numerous transcription factors and co-regulators, as well as histones and the basal transcription machinery are SUMOylated (16). Moreover, genome-wide studies have revealed that SUMOylated proteins are highly enriched at gene regulatory regions, including promoters and enhancers (17–21). Their SUMOylation is likely to occur on chromatin as both SUMO conjugating (E1, E2 and E3s) and deconjugating enzymes can bind to the chromatin (17, 22–24). Although SUMOylation of chromatin-bound proteins has often been associated with gene silencing or gene expression limitation (24–28), it can also participate in the activation of certain genes such as ribosomal genes (19, 20). Overall, the impact of SUMOylation on transcription appears to be dependent on both genes and signaling contexts, as well as on the nature of the conjugated proteins and of the chromatin environment (16).

Here, to better understand the DNA-damaging-independent effects of these drugs, we explored the early effects of Ara-C and DNR on gene expression in AML cells, together with the contribution of SUMOylation to transcriptome reprogramming. We report that DNR induces rapid and broad gene expression changes that are preceded by *de*SUMOylation of chromatin-bound proteins, whereas the effect of Ara-C is much more limited. Intriguingly, we found that *de*SUMOylation limits DNR-induced changes in gene expression. Among the proteins most rapidly *de*SUMOylated in response to DNR, we identified the CTCF insulator protein, which was found highly enriched in regions of the genome marked by SUMO. This notably concerns the *NFKB2* gene, whose DNR-induced expression is preceded by SUMO-dependent rearrangement of chromatin loops involving its SUMO/CTCF-marked promoter and *cis*-regulatory elements.

## Material and methods

### Pharmacologic inhibitors, reagents, and antibodies

Cytosine-β-D-arabinofuranoside (Ara-C), daunorubicin-hydrochloride (DNR), boric acid, protein-G beads, SILAC medium, dimetyl-pimelidade (DMP) were from Sigma. Dialysed serum for SILAC experiments was from Eurobio Abcys. ML-792 was obtained from Takeda Oncology. Lysine and arginine isotopes were from Cambridge Isotope Laboratories. Anti-SUMO-1- (21C7), SUMO2- (8A2) and control- (anti-BrdU, G3G4) hybridomas were obtained from the Developmental Studies Hybridoma Bank (DSHB). The goat polyclonal anti-SUMO-2/3 antibody was described previously (29). The anti-CTCF antibody was from Diagenode (C15410210).

### Cell culture and genotoxic treatment

HL-60 cells were obtained from the ATCC, authenticated by LGC and regularly tested for the absence of mycoplasma. They were cultured at 37°C in the presence of 5% CO_2_ in RPMI (Eurobio) medium supplemented with 10% decomplemented (30 min at 56°C) fetal bovine serum (FBS) and penicillin and streptomycin. After thawing, cells were splited at 0.3 x 10^6^/mL every 2 to 3 days for no more than 10 passages. HEK293T cells were cultured at 37°C in the presence of 5% CO_2_ in DMEM (Eurobio) medium supplemented with 10% decomplemented FBS and penicillin and streptomycin. HL-60 cells were seeded at 0.3 x 10^6^/mL the day before treatment with drugs at 1 µM for DNR and 2 µM for Ara-C. Cells were treated for 2 hrs for ChIP-Seq and 4C experiments and 3 hrs for Affimetrix transcriptomic and RNA-Seq. For SILAC experiments, HL-60 cells were grown in SILAC medium supplemented with dialyzed serum and K0/R0 (light condition), K4/R6 (medium condition), K8/R10 (heavy condition) amino acid isotopes for 21 days until incorporation of amino acids isotopes reached 99%, as measured by mass spectrometry. SILAC labelled cells were then treated or not with 1 µM DNR for 2 hrs. Hybridomas were grown in CellLine bioreactors (Integra) according to the manufacturer’s protocol using RPMI in the cell compartment and RPMI + 10% FCS in the medium compartment. Antibodies were harvested from the cell compartment after 7 days of culture.

### AML patients’ cells

Patient bone marrow aspirates or blood were collected after obtaining written informed consent from patients under the frame of the Declaration of Helsinki and after approval by the Institutional Review Board (Ethical Committee “Sud Méditerranée 1,” ref 2013-A00260-45, HemoDiag collection). Fresh leukocytes were purified using density-based centrifugation using Histopaque 1077 from Sigma and directly lysed for RNA preparation or frozen and stored in liquid nitrogen.

### Gene silencing

The PLK0 lentivirus expressing scramble (SHC002) and UBC9 (NM_003345.3-545S1C1) shRNA expressing vectors were from Sigma. Viral particles were produced and used to transduce HL60 cells as described previously (30). Cells were selected with puromycin (1 µg/mL) for 3 weeks.

### Microarray-based whole transcript expression analysis and profiling

Total RNAs were extracted using the GenEluteTM Mammalian Total RNA kit (Sigma) and treated with DNAse I according to the manufacturer’s specifications. For each condition, 3 independent batches of RNA were prepared and controlled for purity and integrity using the Agilent 2100 Bioanalyzer with RNA 6000 Nano LabChip kits (Agilent Technologies). Only RNA with no sign of contamination or degradation (RIN > 9) were processed to generate amplified and biotinylated sense-strand cDNA targets using the GeneChip® WT PLUS Reagent kit from Affymetrix according to the manufacturer’s specifications. After fragmentation, cDNA targets were used to probe Affymetrix GeneChip® Human Gene 2.0 ST arrays, which were then washed, stained and scanned according to Affymetrix instructions (manual P/N 702731 Rev.3).

### Microarray data analysis

CEL files generated after array scanning were imported into the Partek® Genomics Suite 6.6 (Partek Inc.) for estimating transcript cluster expression levels from raw probe signal intensities using default Partek settings. Resulting expression data were then imported into R (http://www.R-project.org/) for further analysis. First, non-specific filtering was applied to remove transcript clusters with no specified chromosome location. Then, boxplots, density plots, relative log expressions (RLE) and sample pairwise correlations were generated to assess the quality of the data. They revealed no outlier within the series of hybridizations. Principal component analysis (PCA) was also applied to the dataset. The first two components of the PCA could separate samples according to the treatment. Thus, the treatment was considered as the unique source of variability. Finally, the LIMMA package (31) was used to detect differentially expressed genes (DEG) between treated and non-treated samples. A linear model with treatment as unique factor was fitted to the data before applying eBayes function to calculate the significance of the difference in gene expression between the two groups. p-values were adjusted by Benjamin and Hochberg’s False Discovery Rate (FDR) and genes with FDR less than 0.05 and absolute linear Fold Change (FC) greater or equals to 2 were considered as DEG. Microarray data are available at ArrayExpress under the accession number E-MATB-4895.

### RT-qPCR assays

Total mRNAs were purified using the GenElute Mammalian Total RNA kit (Sigma-Aldrich). After 1 hr of DNase I (4U, NEB) treatment in the presence of RNasin (2.5U; Promega), 1 µg of total RNA was used for cDNA synthesis using the Maxima First Strand cDNA kit (Thermo Fisher Scientific). qPCR assays were conducted using Taq platinum (Invitrogen) and the LightCycler 480 device (Roche) with specific DNA primers (Table 1). Data were normalized to the mRNA levels of the housekeeping genes TBP and S26.

**Table 1:**
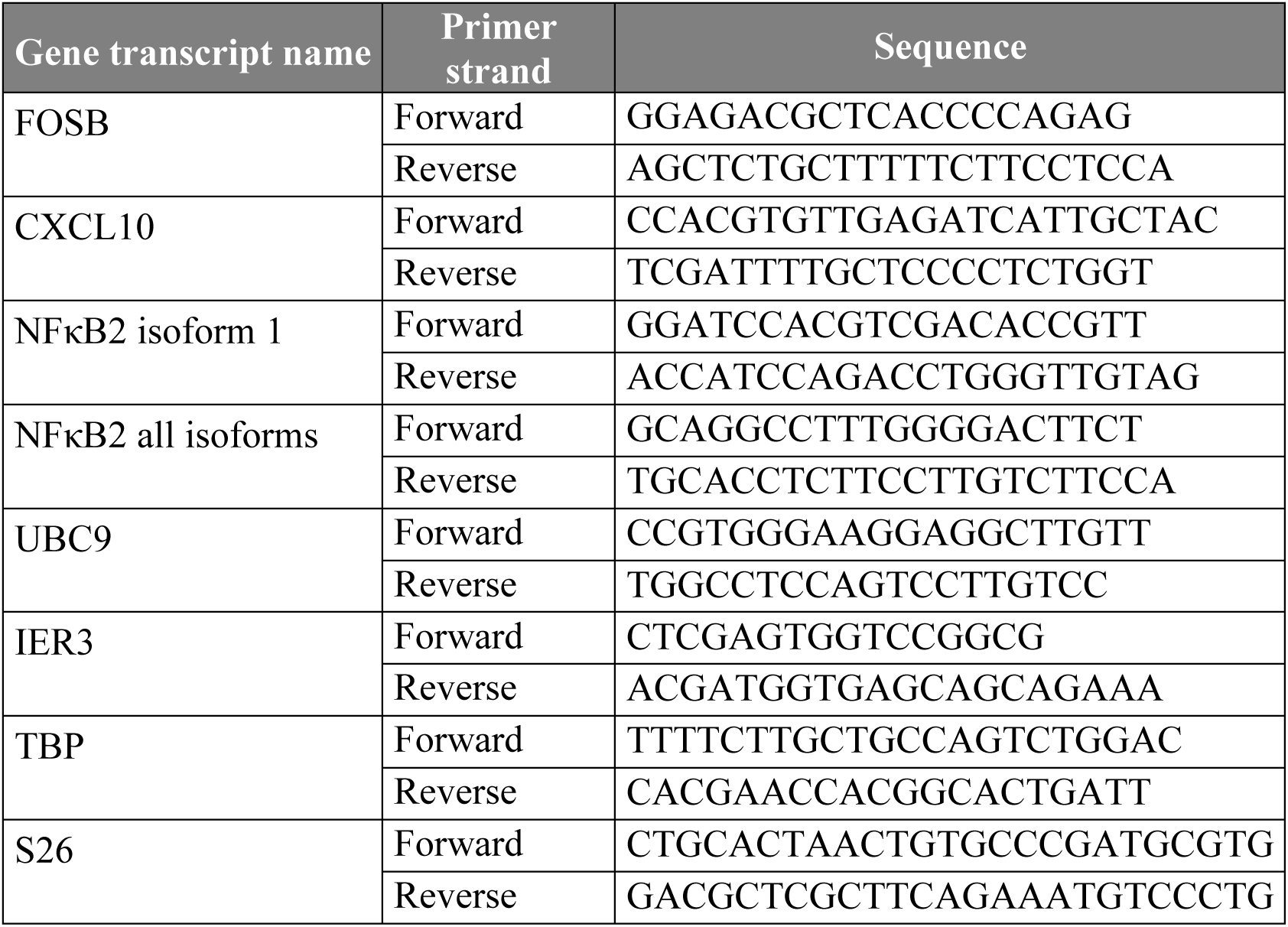
Sequences of the primers used for RT-PCR experiments.

### RNA-seq libraries preparation and sequencing

Total RNAs were purified using the GenElute Mammalian Total RNA kit (Sigma-Aldrich), treated with DNase I (4U; New England Biolabs) in the presence of RNasin (2.5U; Promega) and re-purified. RNA quality was assessed using a BioAnalyzer Nano 6000 chip (Agilent). Three independent experiments were performed. Libraries were prepared using TruSeq®Stranded mRNA Sample Preparation kit (Illumina). Libraries were sequenced using an Illumina Hiseq 2500 sequencer as single-end 50-base reads. Image analysis and base calling were performed using HiSeq Control Software (HCS), Real-Time Analysis (RTA) and bcl2fastq.

### Preparation of DNA for ChIP-seq

A total of 18 x 10^6^ cells were cross-linked with 1% paraformaldehyde for 8 minutes. Paraformaldehyde was then neutralized with 125 mM glycine for 10 minutes. Cross-linked cells were washed with cold PBS, resuspended in a cell lysis buffer (PIPES 5 mM pH7.5, KCl 85 mM, NP40 0.5%, N-ethyl maleimide 20 mM, aprotinin, + pepstatin + leupeptin 1 µg/mL each, AEBSF 1 mM) and incubated at 4°C for 10 minutes. Nuclei were centrifuged (5,000 rpm for 10 minutes at 4°C) and resuspended in a nucleus lysis buffer (Tris-HCl 50 mM pH 7.5, SDS 1%, EDTA 10 mM, N-ethyl maleimide 20 mM, aprotinin + pepstatin + leupeptin 1 µg/mL each, 1 mM AEBSF) and incubated at 4°C for 2.5 hrs. Lysates were then sonicated for 20 cycles of 30 seconds, each at 4°C, using the Bioruptor Pico (Diagenode). After sonication, samples were centrifuged (13,000 rpm at 4°C for 10 min) and the supernatants were diluted 100-fold in the immunoprecipitation buffer (Tris-HCl 50 mM pH 7.5, NaCl 167 mM, N-ethyl maleimide 5 mM, EDTA 1 mM, Triton X100 1.1%, SDS 0.01%, aprotinin + pepstatin + leupeptin 1 µg/mL each, AEBSF 1 mM) with 2 µg of antibodies and Dynabeads Protein G (Thermo Fisher Scientific). Control immunoprecipitation (IP) were performed using the G3G4 antibody (anti BrdU antibody). IPs were performed at 4°C overnight. Beads were then washed in low-salt buffer (Tris-HCl 50 mM pH 7.5, NaCl 150 mM, Triton X100 1%, SDS 0.1%, EDTA 1 mM), high-salt buffer (Tris-HCl 50 mM pH 7.5, NaCl 500 mM, Triton X100 1%, SDS 0.1%, EDTA 1 mM), LiCl salt (Tris-HCl 20 mM pH 7.5, LiCl 250 mM, NP40 1%, deoxycholic acid 1%, EDTA 1 mM), and TE buffer (Tris-HCl 10 mM pH7.5, Tween20 0.2%, EDTA 1 mM). Elution was done in 200 µL of NaHCO3 100 mM containing SDS 1%. Chromatin cross-linking was reversed by overnight incubation at 65°C with NaCl 280 mM followed by 1.5 hrs at 45°C with Tris-HCl 35 mM pH6.8, EDTA 9 mM containing 88 µg/mL of RNAse and 88 µg/mL of proteinase K. Immunoprecipitated DNAs were purified using the NucleoSpin Gel and PCR Clean-up Kit (Macherey-Nagel).

### ChIP-seq libraries preparation and sequencing

For SUMO-2/3 ChIP-seq, immunoprecipitated DNA and corresponding inputs from 3 independent experiments were pooled before library preparation and sequencing. After the analysis of the integrity and the DNA fragment size using the BioAnalyser DNA HS chip (Agilent), ChIP-seq libraries were prepared by the Montpellier MGX platform (https://www.mgx.cnrs.fr) using TruSeq®ChIP Sample Preparation kits (Illumina). The sequencing was processed on Hi-SEQ 2000 (Illumina) as single-end 50 base reads. Image analysis and base calling were performed using HCS and RTA. Demultiplexing was performed using Illumina’s sequencing analysis software (CASAVA 1.8.2) and bcl2fastq.

### 4C-seq experiments

Chromatin for 4C-Seq experiments was prepared essentially as previously described (32, 33). A total of 7 x 10^6^ cells in 10 mL of medium were cross-linked with formaldehyde 2% for 10 minutes at room temperature (RT). Formaldehyde was then neutralized with 125 mM glycine for 10 minutes at 4°C. After a wash with cold PBS, cells were resuspended in 5mL of lysis buffer (Tris-HCl 10 mM pH 8, NaCl 10 mM, NP-40 0.2%, aprotinin + pepstatin + leupeptin 1 µg/mL each, AEBSF 1 mM) and incubated on ice for 20 minutes. Cells were pelleted 5 minutes at 380 g at 4°C, resuspended in 1mL of lysis buffer and snap frozen in liquid nitrogen. Lysates were then thawed at 37°C and centrifuged at 18,000 g at RT for 5 minutes. Cell pellets were resuspended in 700 µL of first enzyme manufacturer buffer 1X (NlaIII – cutsmart [NEB – R0125L]) and homogenized on ice (50 strokes in total) with a 1 mL Dounce homogenizer. Cells were permeabilized using SDS 0.3%, at 37°C for 1 hr under orbital shaking (1 krpm) on an Eppendorf thermomixer). SDS was then displaced by adding TritonX100 1.65% and continuing orbital shaking at 37°C for 1 hr. A 100 µL sample of the reaction mix was taken as a negative control for the first digestion. The digestion with NlaIII enzyme was performed at 37°C for 24 hrs under orbital shaking (1 krpm) using 3 sequential additions of 300 U of enzymes at regular intervals. Before enzyme inactivation at 65°C for 20 min, 100µL of the reaction mix was collected as a restriction enzyme digestion control. The ligation step was performed overnight at 16°C in 8 mL of a reaction mix adjusted to 1X of ligase reaction buffer and containing 800 µL of the restriction enzyme reaction mix, 240 U of T4 DNA ligase HC (Thermo scientific, EL0013) and ATP 0.04mM. Proteinase K (300 µg) was added to ligated DNA products and the reaction was incubated an at 56°C for 1 hr. Decrosslinking was then achieved in an incubation step of 6 hrs at 65°C. The 2 control tubes also underwent the proteinase K and decrosslinking steps. Then, all samples were treated with 300 µg of RNAse at 37°C for 30 minutes. DNA purifications were performed using phenol:chloroform:Isoamyl alcohol 25:24:1 (PCI). DNAs were precipitated at -20°C overnight using 2 volumes of EtOH in the presence of NaCl 250 mM and 20 µg of glycogen (Thermo). DNAs were pelleted by centrifugation (10 krpm) at 4°C and washed using 70% EtOH. Pellets were dried at room temperature and resuspended in 50 µL of water. 10 µL samples were collected from both controls and ligated DNA products and electrophoresed through an agarose gel to control the digestion and ligation steps. Ligation products were digested at 37°C for 2.5 hrs under orbital shaking (1krpm) suing 100U of the second restriction enzyme (DpnII from New England Biolabs, reference R0543M). The second restriction enzyme was inactivated and a second ligation was performed under the same condition as above. 4C libraries were purified with PCI and precipitated as described above. 4C libraries were amplified using specific primers composed of P5/P7 Illumina sequence supplemented with indexes and sequences corresponding to the *NFKB2* promoter (viewpoint) (Table 2). The “Expend Long Template PCR System” kit (Roche) was used using 300 ng of the 4C library following the manufacturer’s instruction. The following amplification parameters were used: denaturation for 2 min at 94°C followed by 30 cycles (94°C - 15 seconds, 58°C - 1 minute and 68°C – 3 minutes) and 7 min at 68°C. 4C libraries were purified with the “Gel and PCR clean up” kit from Macherey-Nagel using NTI solution diluted 6 times and an elution buffer pre-heated at 70°C. After 3 PCR amplification rounds, all 4C libraries for the same sample were pooled, purified and cleaned up using Agencourt AMPure XP beads (ratio 1:1) using EtOH 80% as a washing solution. The libraries were sequenced using the Illumina Hiseq 2500 sequencer as single-end 125 base reads following Illumina’s instructions. Image analysis and base calling were performed using the HiSeq Control Software (HCS), Real-Time Analysis (RTA) and bcl2fastq.

**Table 2:**
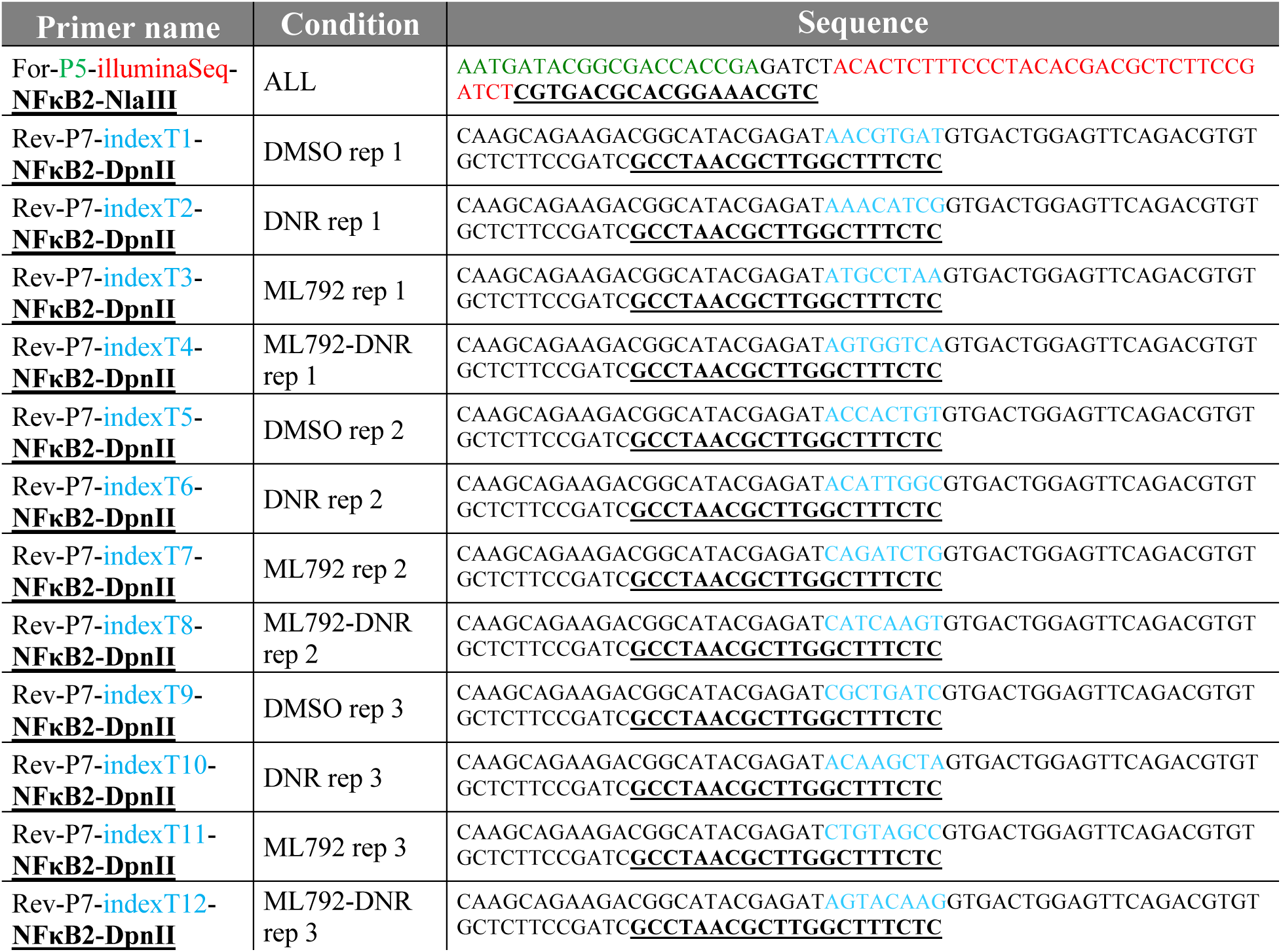
PCR amplification primer to capture NFKB2 promoter interacting regions.

### Quality control of sequencing data and reads trimming

The quality of the data obtained after sequencing was assessed using the FastQC tool. When the score of the first bases of reads was lower than 30, all reads of the dataset were 5’-trimmed of the relevant number of nucleotides using the trimmomatic tool (Headcrop). All reads with more than 1 N-call were removed from datasets.

### ChIP-seq reads mapping, peak calling and analysis

ChIP-seq reads were aligned on the human reference genome (hg19) using CASAVA 1.8.2 (MGX pipeline). Analysis of the aligned reads, scaling and input subtraction were performed using the R package Pasha (34). Data were visualized using the IGB software (35). The peak calling was performed using the WigPeakCaller script, which automatizes the IGB thresholding tool (36). The SUMO-2/3 peak calling was done with the following parameters: by value=32, Max Gap ≤ 100 and Min Run > 100. Motif search was performed using HOMER v4.10 (37). ChIP-Seq sequencing data are available with accession GSE198986. Publicly available HL-60 ChIP-seq dataset were used for CTCF (GSM749688), H3K4me3 (GSM945222), H3K4me1 (GSM2836484), H3K27ac (GSM2836486) and RNAPII (GSM1010737). The hg19 promoter (−2kb to TSS) gff files have been generated with gff_toolbox, using the GRCh37p13 annotation file from NCBI. The H3K4me3 histone marks, which is enriched at gene TSS, have been used as a proxy to annotate HL-60 promoter. All genomic regions presenting H3K4me1, which do not correspond to annotated promoters, were considered as candidate enhancers. Then, the activity of these regulatory elements was inferred from the presence of H3K27ac. All dataset intersects were performed using Bedtools 2.29.0 (intersect) from Quinlan laboratory (38, 39).

### RNA-seq mapping, quantification and differential analysis

RNA-seq reads were mapped to Human reference genome (hg19, GRCh37p13) using TopHat2 (2.1.1) (40) based on the Bowtie2 (2.3.5.1) aligner (41). The reproducibility of replicates was quantified using the cufflinks v2.2.1 tool (42) with the linear regression of reads per kilobase million (RPKM) between 2 replicates. Read association with annotated gene regions was done using the HTseq-count tool v0.11.1 (43). The variance between replicates and conditions were appreciated thanks to a principal component analysis (PCA) performed on the read count matrix. Differential expression analysis was performed using DESeq2 (44) using the normalization by the sequencing depth and the parametric negative binomial law to estimate data dispersion. All conditions were compared to the mock condition (DNR *vs* DMSO, ML-792 *vs* DMSO and ML-792+DNR *vs* DMSO) and the ML-792+DNR condition was also compared to the DNR-only condition (ML-792+DNR *vs* DNR). The genes that presented a fold change ≥ or ≤ 2 and an adjusted p-value (FDR) < 0.05 were considered as differentially expressed genes (DEGs). RNA-seq data are available with accession GSE198986.

### 4C-seq mapping, trim, capture and profiling

The pipeline for the analysis of the 4C data was modified from the pipe4C pipeline (45) and is available on github (https://github.com/Mathias-Boulanger/pipe4C). The steps are the following: Reads filtering (trim-capture), mapping to reference genome, assignment of reads to their restriction fragment and creation of normalized score per fragment. Only reads containing the amplification sequence (CGTGACGCACGGAAACGTC) were kept for further analysis. Then, sequences downstream of the restriction enzyme cutting site of each selected reads were mapped to GRCH37p13 human reference genome with Bowtie2 aligner. Restriction fragment map was extrapolated from the reference genome using the cutting sequence of restriction enzymes. The interaction peak calling has been performed with peakC and the differential profiling analysis with DESeq2 (44, 46). 4C-seq data are available with accession GSE198986.

### Gene Ontology

Functional gene-annotation enrichment analysis were done using GO Panther (47) with the ID number of DEGs or proteins as input list. The gene network analyses were performed using the Cytoscape-based Cluego plugin (48).

### Coupling antibodies to protein-G beads

Hybridoma supernatants were incubated with Protein G sepharose beads (SIGMA) at room temperature for 4 hrs, washed 3 times with PBS (phosphate buffer 10 mM pH 7.4, KCl 2.7 mM and NaCl 137 mM) and once with Na borate 50 mM pH 9.0. Antibodies were then crosslinked for 30 min in dimethyl-pimelimidate (DMP) 20 mM diluted extemporarily in Na borate 50 mM pH 9.0. The coupling procedure was repeated a second time and the beads were washed 3 times with PBS.

### Immunoprecipitation of SUMOylated proteins

For SILAC experiments, SILAC-labeled HL-60 cells were grown in spinner flasks (Nunc). 5 x 10^8^ cells were used for each condition. The immunoprecipitation of endogenously SUMOylated proteins was based on the protocol described in reference (49). Cells were lysed in PBS containing SDS 2%. The final concentration of SDS after lysis was then adjusted to 1% and lysates were sonicated. Dithiotreitol (DTT) was then added at a final concentration of 50 mM. Lysates were then boiled for 10 min and diluted 10-fold in Na phosphate 20 mM pH 7.4, 150 mM NaCl, Triton X100 1%, Na deoxycholate 0.5%, EGTA 5 mM, EDTA 5 mM, NEM, 10 mM, aprotinin + pepstatin + leupeptin 1 µg/mL each, filtered through 0.45 µm filter and incubated with Protein G-coupled anti-SUMO-1, -SUMO-2 and -BrdU (control) antibodies at 4°C overnight. Beads were then washed 3 times with RIPA (Na phosphate 20 mM pH 7.4, NaCl 150 mM Triton X100 1%, SDS 0.1%, Na deoxycholate 0.5%, EGTA 5 mM, EDTA 5 mM, NEM 10 mM, aprotinin1µg/mL and pepstatin 1 µg/mL) and twice with RIPA containing NaCl 350 mM in LowBind tubes (Eppendorf). Elution of SUMOylated proteins was performed twice with peptides bearing either the 21C7 SUMO-1- (VPMNSLRFLFE) or the 8A2 SUMO-2/3- (IRFRFDGQPI) epitope diluted in RIPA containing NaCl 350 mM. Eluted proteins were precipitated with 10% TCA for 1 h on ice. Pellets were then washed twice with acetone at - 20°C, dried and resuspended in the Laemli electrophoresis sample buffer. For the identification of SUMOylated targets (SILAC1), samples were immunoprecipited with control-, anti-SUMO-1 or anti-SUMO-2/3 antibodies and mixed only after elution with the SUMO epitope-bearing peptides. For the identification of proteins showing DNR-modulated SUMOylation, mock- and DNR-treated samples were mixed right after the initial lysis step and used for immunoprecipitation with SUMO-1 (SILAC2) or SUMO-2/3 (SILAC3) antibodies.

### Mass spectrometry identification of SUMOylated proteins

Enriched SUMOylated proteins from SILAC lysates were size-separated by SDS-PAGE and in-gel digested with trypsin. The resulting peptide mixtures were extracted, desalted and concentrated on STAGE-tips with two C18 filters and eluted two times with 10 μl of acetonitrile 40% in formic acid 0.5% prior to online nanoflow liquid chromatography-tandem mass spectrometry (nano LC-MS/MS) using an EASY-nLC system (Proxeon, Odense, Denmark) connected to the Q Exactive HF (Thermo Fisher Scientific, Germany) through a nano-electrospray ion source. Peptides were separated in a 15 cm analytical column in-house packed with 1.9 μm C18 beads (Reprosil-AQ, Pur, Dr. Manish, Ammerbuch-Entringen, Germany) using an 80 minutes gradient from 8% to 75% acetonitrile in acetic acid 0.5% at a flow rate of 250 nl/minute. The mass spectrometers were operated in data-dependent acquisition mode with a top 10 method. For Q-Exactive measurements, full scan MS spectra were acquired at a target value of 3 x 106 and a resolution of 60,000 and the Higher-Collisional Dissociation (HCD) tandem mass spectra (MS/MS) were recorded at a target value of 1 x 105 and with a resolution of 60,000 with a normalized collision energy of 30%.

Raw mass spectrometry (MS) files were processed with the MaxQuant software suite (version 1.4.0.3, www.maxquant.org). All resulting MS/MS spectra were searched against the human Uniprot database (www.uniprot.org) by the Andromeda search engine using the reversed database strategy applying a false discovery rate of 0.01 at both peptide and protein levels. Overrepresentation of Gene Ontologies of the identified proteins were analyzed using Fisher’s exact test from InnateDB (50).

### Statistical analyses

Results are expressed as means ± S.D. Statistical analyses were performed using the paired Student’s *t*-test with the Prism 5 software. Differences were considered as significant for P-values of <0.05. *; **; *** correspond to p < 0.05; p < 0.01; p < 0.001, respectively. ns=not significant. Statistical analyses of the transcriptomic and proteomic experiments are described in the relevant sections.

## Results

### DNR and Ara-C rapidly induce transcriptional programs related to cell proliferation/death- and inflammation/immunity in AML cells

To identify the genes whose expression is rapidly altered by Ara-C or DNR in AML cells, we performed a whole transcriptome profiling of HL-60 cells, one of the most widely used cellular model of AML (51). Cells were treated with each one of the two drugs at doses relevant to the clinical practice (2 and 1 µM, respectively) (52, 53) for 3 hrs, *i.e*. before the onset of apoptosis, which begins after 4 hrs of treatment (13). Using the Affimetrix array technology, we identified 476 significant differentially expressed genes (DEGs) in DNR-treated cells, 182 being upregulated and 294 downregulated more than 2-fold (Figure 1A and Supplementary Table 1). Much less DEGs were identified in Ara-C-treated cells: 6 were upregulated and 29 downregulated by a >2-fold factor (Figure 1B). Gene ontology (GO) enrichment analyses revealed that the genes identified as down-regulated upon treatment by Ara-C and/or DNR are mostly involved in nucleosome assembly (Supplementary Figure 1A). Those up-regulated principally belong to functional categories linked to signal transduction, transcription, cell proliferation and death (with both pro- and anti-apoptotic genes being induced) and inflammation/immunity (Figure 1C, 1D, Supplementary Figure 1B, Supplementary Table 1). We confirmed the activation of four of the most DNR-induced genes (*CXCL10*, *FOSB*, *NFKB2* and *IER3*) by RT-qPCR in HL-60 cells treated with DNR (Figure 1E). Noteworthy, these genes were not significantly induced by Ara-C even at concentrations higher than 2 µM (up to 50 µM) (Figure 1E). Taken with our Affymetrix data (Figure 1A and 1B), this suggested that DNR is more potent at altering transcription than Ara-C in the HL-60 cell model. We then analyzed samples from three AML patients taken at diagnosis. These were treated *ex vivo* with DNR or Ara-C for 3 hrs and assayed for the expression of the same four genes. All of them were induced by DNR in the three patients tested, albeit to different degrees. Their expression was more induced by DNR than by Ara-C for two patients, showing that our observation in HL60 cells reflected a situation happening in primary AML cells. However, the reverse was observed for the third patient sample, which is probably reflecting AML heterogeneity (Figure 1F).

**Figure 1:**
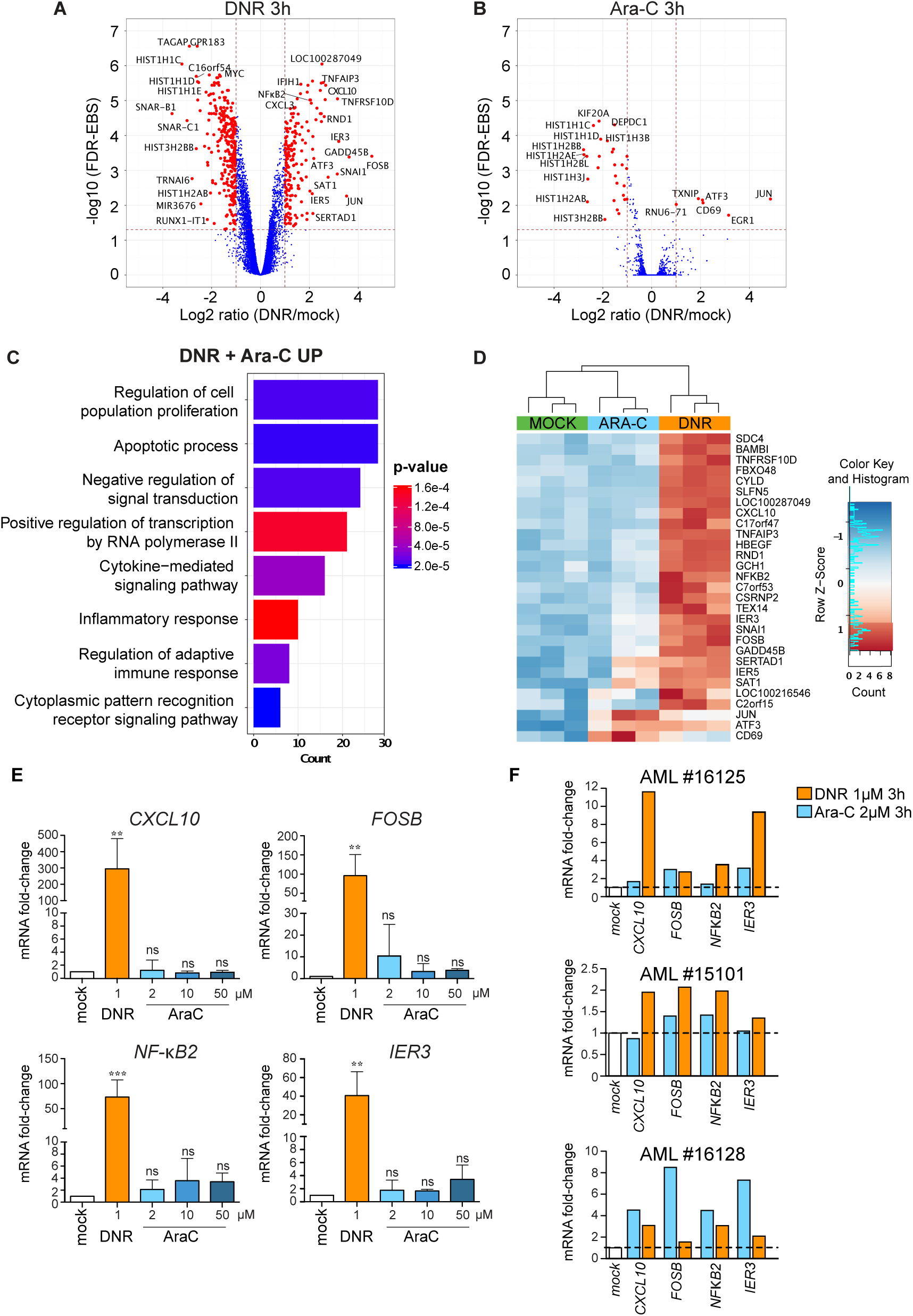
Chemotherapeutic drugs rapidly alter the expression of genes involved in cell death and inflammation in AML cells. *A-B: trancriptome profile.* HL-60 cells were treated with 1 µM DNR (A) or 2 µM Ara-C for 3 hours (B). RNAs were purified from 3 independent experiments and used to probe Affymetrix Human Gene 2.0 ST Genechips. The red dots on the Volcano plots represent the Differentially Expressed genes (DEG) with an absolute Fold Change (FC) ≥ 2 (log2 ≥ 1) and a False Discovery Rate (FDR) corrected with Empirical Bayes Statistics (EBS)(76) < 0.05. *C: Gene Ontology enrichment analysis of the genes up-regulated (*≥ *2 fold) by DNR and Ara-C.* Ontologies were performed using the Panther GO database (47). The main terms of each identified group are presented on the graph and classified by the number of genes present in each group. *p* values are corrected with Bonferroni step down. *D: Heatmap of DEG with a FC* ≥ *4 in the transcriptomic experiments presented in A and B*. The data for all 3 replicates are represented. *E: RT-qPCR analysis of selected genes*. HL-60 cells were treated for 3 hours with 1 µM DNR or 2 µM Ara-C. The levels of the indicated mRNAs were measured by RT-qPCR, normalized to *GAPDH* levels and expressed as fold increase to mock-treated cells (mean +/- SD, n=7 for *NF-κB2*, n=6 for *IER3*, n=5 for *FOSB*, *CXCL10*). *F: Regulation of selected genes in primary AML cells.* AML cells (bone marrow aspirate) from 3 patients were treated *in vitro* with 1 µM DNR or 2 µM Ara-C for 3 hours. The levels of the indicated mRNAs were measured by RT-qPCR, normalized to *TBP* levels and expressed as fold increase to mock-treated cells.

Thus, our data indicate that one early effect of the chemotherapeutics used as frontline treatment of AML is transcriptional reprogramming. DNR, however, shows broader effects than Ara-C and the genes most induced by DNR principally belong to two general functional categories: cell proliferation/death and inflammation/immunity.

### DNR induces a fast removal of SUMO from chromatin

We have previously shown that DNR and Ara-C induce a progressive *de*SUMOylation of cellular proteins in AML. It starts around 2 hrs after the beginning of the treatment and is due to the inactivation of the SUMO E1 and E2 enzymes via the formation of a ROS-dependent disulfide bond between their catalytic cysteines (13). Due to the role of SUMO in transcription, we wondered whether DNR and Ara-C could induce fast alterations in SUMOylated protein distribution on the genome, as such changes might contribute to drug-induced transcriptional changes. This was addressed in ChIP-Seq experiments with antibodies directed to SUMO-2/3 as these paralogs are more enriched on chromatin than SUMO-1 (25). HL-60 cells were treated with DNR or Ara-C for 2 hrs, *i.e*., a time point earlier than that used in our transcriptomic analysis to consider the time required between gene transcription alteration and RNA accumulation changes in the cell. In untreated cells, and as previously shown by others (17, 18, 26, 54, 55), SUMO-2/3 was found distributed all along chromatin with approximately 44,000 peaks. A particular enrichment was found at both annotated gene promoters and candidate enhancer regions defined by the presence of high H3K27ac, H3K4me1 and low H3K4me3 (Figure 2Aa and Supplementary Figure 2). In mock-treated cells, we identified 6861 genes showing a significant accumulation of SUMOylated proteins in their promoter regions with a peak of enrichment approximately 100 bp upstream of Transcription Start Sites (TSSs) (Figure 2B). Interestingly, SUMOylated proteins were found enriched on active promoter regions (those with high H3K4me3 and RNApolII) and not on inactive ones (those with low H3K4me3 and RNAPolII) (Supplementary Figure 2A). Along the same line, SUMOylated proteins were found localized in the center of the candidate enhancer regions (Figure 2C) and slightly more enriched on active- (*i.e.* with high H3K27ac) than on inactive- (*i.e.* with low H3K27ac) candidate enhancers (Supplementary Figure 2B).

**Figure 2:**
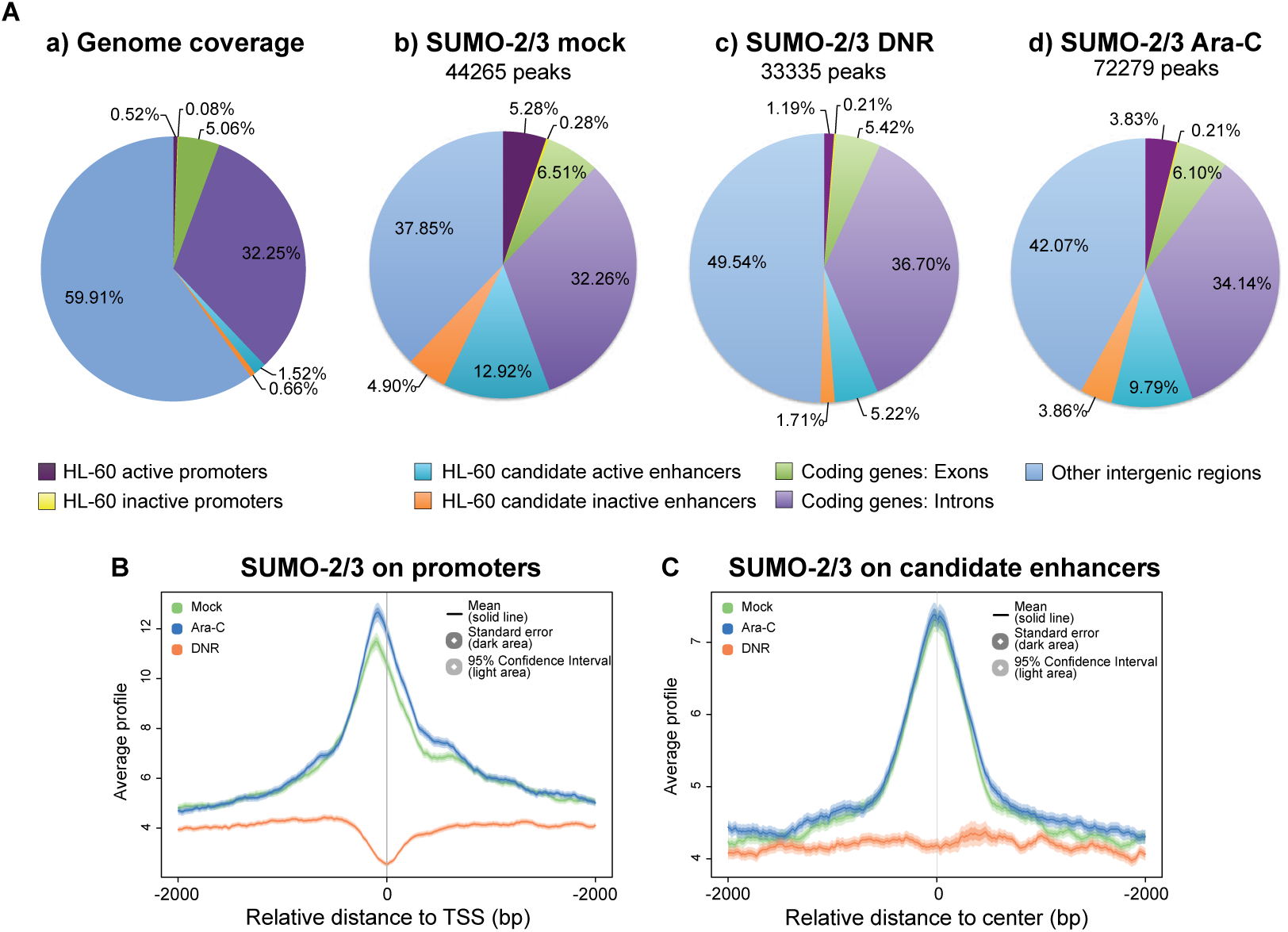
Treatment of AML cells with DNR depletes SUMOylated proteins from the chromatin, in particular at promoters and enhancers. *A and B: ChIP-Seq analyses of SUMO-2/3 distribution on the genome.* HL-60 were treated with 1 µM DNR or 2µM Ara-C for 2 hours. ChIP-Seq experiments were carried out with SUMO-2/3 antibodies. *A*: Distribution of SUMO-2/3 on chromatin in mock-, DNR- or Ara-C-treated HL-60 cells. *B and C:* Metaprofile of the SUMO-2/3 ChIP-seq signal on HL-60 promoter (B) or enhancer (C) in mock-, DNR- or Ara-C-treated HL-60 cells. Promoters (−2kb to TSS) and enhancers as well as their activation state were defined using H3K27ac, H3K4me1 and H3K4me1 profiles as well as NCBI refseq data (see Material and methods and Supplementary Figure 2).

We then analyzed whether DNR and Ara-C treatments globally affected the presence and/or the distribution of SUMO-2/3-conjugated proteins on chromatin. Surprisingly, the Ara-C treatment, which moderately modulated gene expression in HL-60 cells (Figure 1B), led to an increased number of SUMO peaks on the genome (Figure 2A). However, their overall distribution amongst the different regions of the genome remained essentially unchanged as compared to untreated cells (Figure 2A) and the same held true for their distribution at the level of promoters (Figure 2B) and enhancers (Figure 2C). By contrast, the DNR treatment, which was more potent at altering gene expression in HL-60 cells, induced a 25% decrease in the total number of SUMO peaks. This decrease in the presence of SUMOylated proteins on chromatin was much stronger at promoters (Figure 2B) and enhancers (Figure 2C). As mentioned earlier, although *de*SUMOylation starts after 2 hrs of DNR treatment as seen by the accumulation of free SUMO, the bulk of protein SUMOylation is not detectably affected at this early time point (13). This raises the idea that proteins bound to gene *cis*-regulatory regions are among the first cellular proteins to be *de*SUMOylated upon DNR treatment.

### Inhibition of SUMOylation limits both positive and negative DNR-induced transcriptome changes

As DNR had much stronger effects on chromatin SUMOylation and gene expression than Ara-C, we continued our investigations by assessing whether inhibition of SUMOylation is sufficient to induce the expression of DNR-responsive genes. To this aim, we performed RNA-seq analyses of HL-60 cells treated for 3 hrs with the highly potent and selective SUMOylation inhibitor ML-792 (56). We found that ML-792 had minimal effect on gene expression with only 21 differentially regulated genes (Figure 3A), suggesting that *de*SUMOylation *per se* is not sufficient to induce DNR-responsive genes. As there is no specific *de*SUMOylation inhibitors that could be used to prevent DNR-induced *de*SUMOylation, we used ML-792 in combination with DNR to amplify/accelerate DNR-induced deSUMOylation. RNA-Seq being more sensitive than the Affimetrix array-based approach, we identified more DNR-responsive genes than in our former transcriptomic approach (Figure 1). 552 genes were found up-regulated and 380 down regulated in DNR *vs* mock-treated cells (Figure 3A and Supplementary Table 2). ML-792 did not strongly affect the number of DNR-regulated genes, as 521 of them were still found up-regulated and 337 down-regulated (Figure 3C). However, the comparison of ML-792+DNR- to DNR only-treated-cells revealed that inhibition of SUMOylation during the DNR treatment limited their up- or down-regulation. This was in particular the case for the genes, which are the most affected by DNR (Figure 3D and 3E). GSEA analysis showed that all pathways enriched in DNR-treated cells were less enriched when SUMOylation was inhibited, the most pronounced effects being observed for the genes involved in inflammation (Figure 3F). Thus, our data suggest that rapid DNR-induced removal of SUMO from chromatin counteracts the ability of DNR to alter the expression of the vast majority of its responsive genes, whether induced or down-regulated.

**Figure 3:**
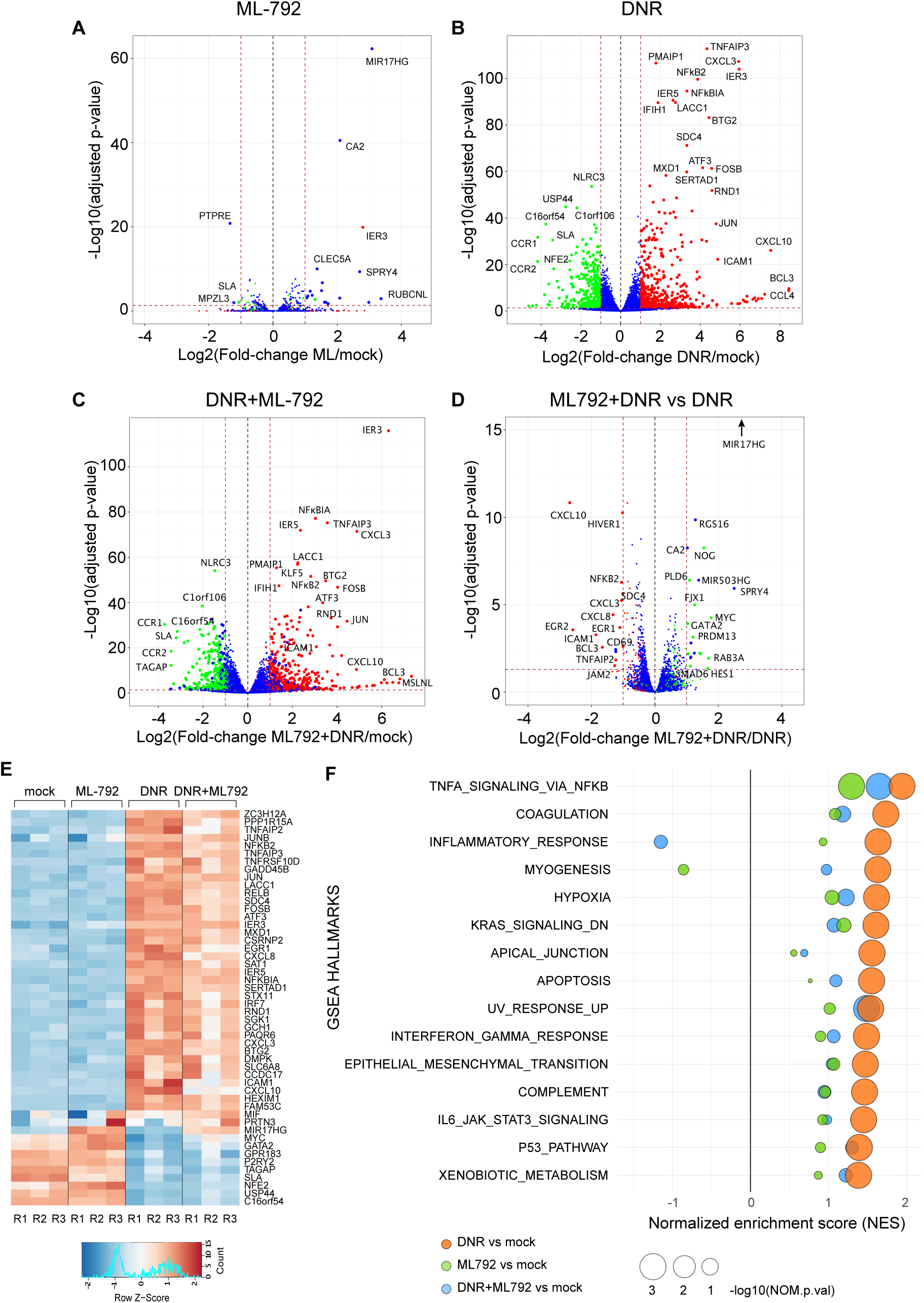
Inhibition of SUMOylation reduces the DNR-induced regulation of a subset of genes. (A-E) HL-60 cells were treated with 1 µM DNR, 0.5 µM ML-792 or the combination of the two drugs for 3 hrs. Total RNAs were prepared from 3 independent experiments and sequenced. Volcano plot showing the DEG between (A) ML-792- and mock- (B) DNR- and mock-, (C) ML-792+DNR- and mock-, (D) ML-792+DNR and DNR-treated HL-60 cells. Green dot: DNR-downregulated FC ≤ -2 and FDR (false discovery rate) < 0.05; Red dots: DNR-upregulated with FC ≥ 2 and FDR < 0.05; Blue dots: genes with -2 ≥ FC ≤ 2 and FDR > 0.05 in the DNR *vs* mock conditions. (E) Heatmap of top 50 DEGs in all conditions presented in A, B and C. (F) Gene Set Enrichment Analysis (GSEA) was performed using RNA-Seq data presented in A-E. The GSEA hallmarks showing a Normalized Enrichment Score NES>1 or <-1, a p-value<0.05 and an FDR <0.25 for the DNR *vs* mock analysis are presented for each treatment condition (DNR, ML-792, DNR+ML-792) compared to the mock-treated cells.

### Transcriptional factors and co-regulators are the fastest and main class of deSUMOylated proteins upon DNR treatment

To better understand how *de*SUMOylation controls DNR-responsive gene expression, we next resorted to large-scale proteomics to identify the proteins changing their SUMOylation levels after 2 hrs of DNR treatment, *i.e.*, the time point at which important changes in chromatin protein SUMOylation were detected by ChIP-seq (Figure 2). First, we characterized the HL-60 cell proteome conjugated to SUMO-2/3 but also to SUMO-1. To this aim, we immunoprecipitated and identified by quantitative mass spectrometry SUMO-2/3- and SUMO-1 modified proteins. 894 SUMO targets were identified, most of them being modified by both SUMO-2/3 and SUMO-1 (Figure 4A). Then, SUMO-2/3 or SUMO-1-conjugated proteins were immunoprecipitated and identified from HL-60 cells treated or not with DNR for 2 hrs. The SUMOylation level of most proteins did not change after this short treatment with DNR. However, 34 proteins (31 for SUMO-2/3 and 11 for SUMO-1, 8 proteins being common) showed increased modification (Figure 4B and Supplementary Table 3). Much more proteins (83 for SUMO-2/3 and 32 for SUMO-1, 19 being common) showed a significant decrease in their SUMO conjugation after DNR treatment (Figure 4B and Supplementary Table 3). Finally, these changes were not due to modifications of protein abundance, as determined by sequencing of input samples in control- and DNR-treated cells (Supplementary Table 3). Interestingly, after 2 hrs of treatment, most of the *de*SUMOylated proteins (both SUMO-2/3 and SUMO-1 substrates) were found to be chromatin-bound proteins involved in the regulation of gene expression (Figure 4C).

**Figure 4:**
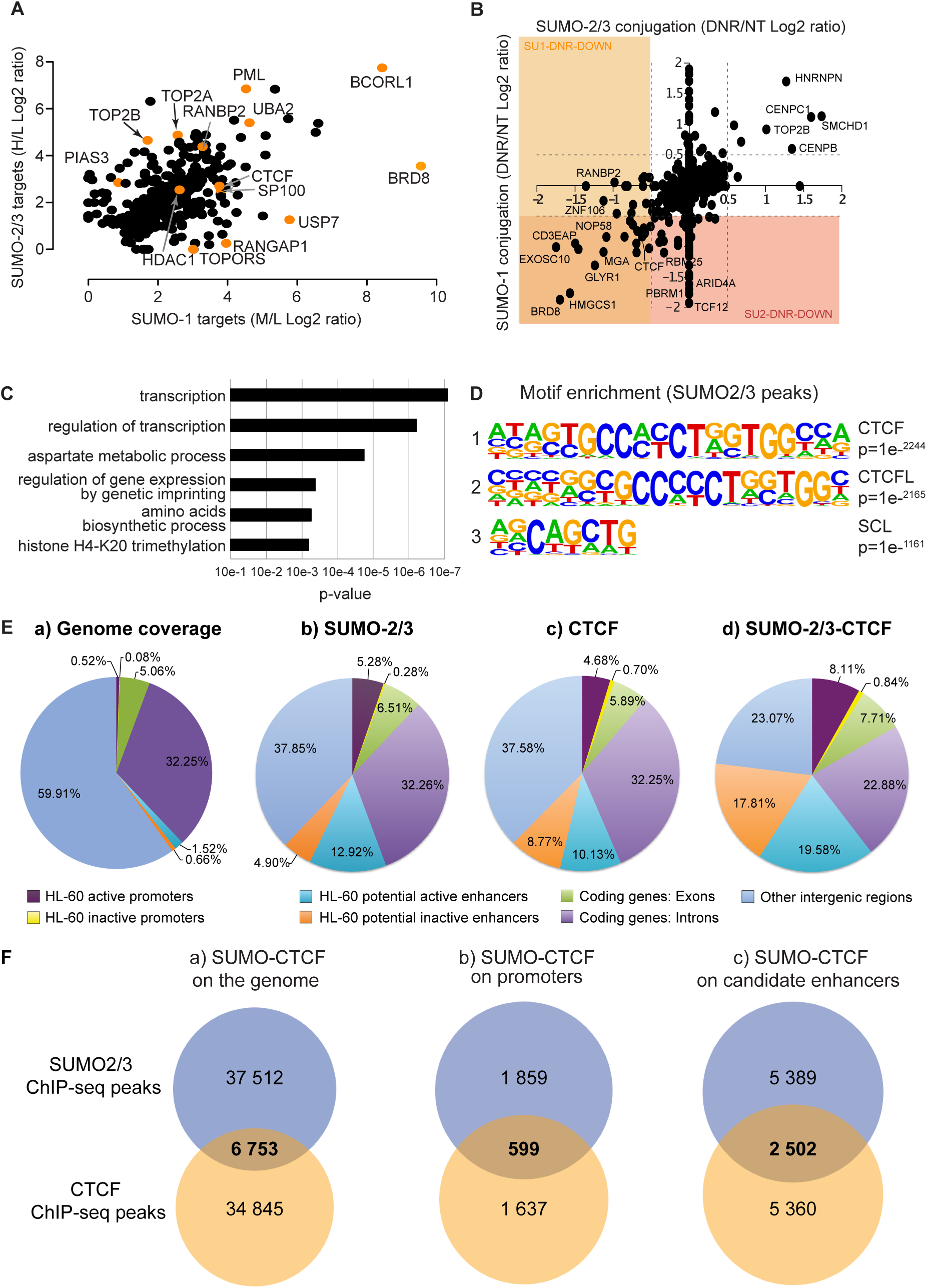
DNR leads to *de*SUMOylation of chromatin regulators, including CTCF. *A: Endogenously SUMOylated proteins in HL-60 cells.* SUMOylated proteins were immunoprecipitated from triple SILAC-labeled HL-60 cells with control (light condition), anti SUMO-1 (medium condition) or anti SUMO-2/3 (heavy condition) antibodies and identified by mass spectrometry. Log2-transformed ratios between SUMO-1 or SUMO-2/3 and control IP are represented. Only proteins with more than 2 peptides identified and a Log2 ratio>1 (SUMO-1/control or SUMO-2/control) were selected. The orange dots correspond to selected known SUMO substrates. *B: Changes in SUMO-1 and SUMO-2/3 proteomes upon DNR treatment.* Scatterplot analysis of SUMO-1 and SUMO-2/3 proteome change (Log2 ratio) in cell treated with DNR (1 µM for 2 hrs) compared to mock-treated cells. Doted lines represent Log2 ratio of -/+ 0.5. Only proteins found to be SUMOylated in A are represented. C: *DeSUMOylated proteins are mostly transcriptional regulators*. Gene Ontology analysis of the identified down-SUMOylated proteins for SUMO-1 and SUMO-2/3 in response to DNR (log2 ratio <-0.5) were obtained using the Panther Protein Class database (47). *D: The CTCF motif is enriched at SUMO-2/3 binding sites.* Motif enrichment search was performed with homer pearl script (findMotifs.pl) on the SUMO-2/3 ChIP-Seq data obtained for mock-treated HL-60. The 3 most enriched motifs are shown. *E: SUMO/CTCF overlap on promoter and enhancers.* Venn Diagrams for the distribution of the sequences bound by (b) SUMO-2/3, (c) CTCF and (d) both proteins. *F: Venn Diagram for the distribution of SUMO-2/3 and CTCF* peaks on the whole genome, on promoters (−2 kb;TSS) and putative enhancers in HL-60 cells.

Thus, our proteomic data support the idea initially raised by our SUMO-2/3 ChIP-seq experiments (Figure 3) that chromatin-bound proteins are among the first to be *de*SUMOylated upon treatment by DNR. As *de*SUMOylated proteins are principally involved in gene expression regulation, this further suggests that the SUMO pathway regulates the expression of DNR-responsive genes in AML cells.

### CTCF binds to promoters of the majority of DNR-responsive genes and is *de*SUMOylated upon DNR treatment

Among the SUMO substrates that are rapidly *de*SUMOylated upon DNR treatment (Figure 4B), we noted the CCCTC-binding factor CTCF, an insulator protein known to regulate the three-dimensional architecture of chromatin (57). CTCF was formerly reported to be SUMOylatable (58) and its SUMOylation to be instrumental for activation and repression of the *PAX6* (59) and *c-MYC* (60) genes, respectively. We therefore considered that its fast *de*SUMOylation in response to DNR might contribute to the changes in DNR-responsive gene expression in AML cells. As a first indication supporting this hypothesis, we found that the DNA-binding motif most represented under the SUMO peaks identified in our ChIP-seq experiments (Figure 3) was the consensus CTCF-binding motif (Figure 4D). Interestingly, when analyzing publicly available CTFC ChIP-seq data obtained in HL-60 cells (61), we found that CTCF, like SUMO-2/3, is strongly enriched at gene *cis*-regulatory elements (*i.e.* promoters and enhancers; Figure 4Ea). We then addressed the colocalization of CTCF and SUMO in HL-60 cells by crossing their respective ChIP-seq data. At the genome-wide level, 15% (6,753) of the SUMO-2/3 peaks overlapped with CTCF ones (Figure 4Fa). Of these overlapping peaks, nearly half (46%) were found at either promoters or candidate enhancers (Figure 4Ed), which is twice the percentage of SUMO-2/3 (23.38%) or CTCF (24,28%) peaks present at the same loci. This indicated that CTCF/SUMO-2/3 bound sites are mostly found in gene *cis*-regulatory regions. In total, overlapping SUMO-2/3 and CTCF peaks were found at 599 gene promoters and 2502 candidate enhancers (Figure 4Fb and 4Fc), suggesting a functional interplay between SUMO and CTCF at, at least a fraction of, these genetic elements in HL-60 cells.

Finally, we asked to which extent SUMO levels were affected upon treatment by DNR or Ara-C at the CTCF-bound sites marked by SUMO. To this aim, we crossed CTCF ChIP-seq data with our SUMO-2/3 ChIP-seq data obtained in mock-, DNR- and Ara-C-treated HL60 cells. The only SUMO peaks that were not affected by Ara-C were those co-localizing with CTCF signals (Supplementary Figure 3A). Considering that the number of total SUMO-2/3 peaks increases after Ara-C treatment (Figure 2), this suggested that the SUMOylated proteins that are present at CTCF-bound sites are protected from *de*SUMOylation and/or re-localization to other genomic regions upon Ara-C treatment. In contrast, DNR, which affects gene transcription to a much greater extent than Ara-C, led to a complete depletion of SUMO at the loci co-marked by SUMO and CTCF (Supplementary Figure 3A). Strengthening the finding of massive loss of SUMO-2/3 at CTCF-binding sites in the presence of DNR, but not in that of Ara-C, Pearson’s correlation coefficients of CTCF- *vs* SUMO-2/3 signals at promoters and enhancers were comparable in mock- and Ara-C-treated HL-60 cells, whereas they were dramatically diminished upon DNR treatment (Supplementary Figure 3B, 3C).

Thus, our data indicate that Ara-C, which has minimal effects on gene expression, does not detectably affect SUMOylation at CTCF-bound loci, whereas DNR, which entails larger transcriptomic changes, induces strong *de*SUMOylation of the SUMO-marked gene regulatory elements bound by CTCF. This raises the idea that *de*SUMOylation of CTCF itself and/or of proteins associating/colocalizing with CTCF at promoters and/or enhancers regulates the expression of DNR-responsive genes.

### SUMOylation regulates DNR-induced expression of the CTCF- and SUMO-bound *NFKB2* gene

To further investigate the links between CTCF and SUMO in DNR-induced gene expression changes, we crossed the list of genes presenting SUMOylated proteins and CTCF at the same position in their promoters with that of genes transcriptionally regulated by DNR. Sixty-seven genes were identified, the expression of which might be regulated through SUMOylation/*de*SUMOylation of proteins (including CTCF) bound to their promoter regions (Figure 5A, left panel). We then crossed this list of 67 genes with that of the 36 genes whose DNR-induced expression changes were altered by more than 2-fold upon SUMOylation inhibition (Figure 5A, right panel). This led to the identification of 4 genes (*EGR1*, *ICAM1*, *MYC* and *NFKB2*) whose DNR-induced up- or down-regulation is reduced by the inhibition of SUMOylation and whose proximal promoters are marked by SUMO and CTCF at the same positions. We then focused on the *NFKB2* gene, encoding the transcription factor Nuclear Factor-kappa B2 (NF-κB2), because of its involvement in the regulation of both cell death/survival and inflammation/immunity (62, 63), processes we found associated with the response of AML to DNR. Moreover, after having formerly shown that DNR induces *NFKB2* expression in AML patients’ cells treated *in vitro* (Figure 1F), we established the early induction of this gene *in vivo* using peripheral blood mononuclear cells (PBMCs) purified from 3 AML patients before and 4 hours after the beginning of an induction chemotherapy comprising DNR and Ara-C (Figure 5B). We also confirmed our RNA-Seq results showing that *NFKB2* is strongly induced by DNR in HL-60 cells using a RT-qPCR. We found a 16-fold induction when all *NFKB2* mRNA isoforms were analyzed together and a 70-fold induction when considering only its longest isoform, which starts at the CTCF/SUMO bound site (Figure 5C). Moreover, consistent with our RNA-Seq data (Figure 3), the SUMOylation inhibitor ML-792 strongly decreased the DNR-induced expression of *NFKB2* (Figure 5C). Importantly, ML-792 also prevented the induction of *NFKB2* by DNR in primary AML cells treated *ex vivo* (Figure 5D). To further confirm the implication of SUMOylation inhibition in this process, we resorted to RNAi to down-regulate the SUMO E2 enzyme Ubc9. This did not affect the basal level of *NFKB2* expression but limited its DNR-induced up-regulation (Figure 5E). ChIP-Seq data identified a major SUMO-2/3 peak (Peak 1) colocalizing with CTCF at the most 5’ promoter of *NFKB2* in HL-60 cells, as well as a second CTCF-SUMO peak (Peak 2) in the gene body (Figure 5F). Both peaks disappeared after DNR treatment and, consistent with the limited effect of Ara-C on *NFKB2* expression, were much less affected by the Ara-C treatment (Figure 5F). Thus, altogether, our results suggest that *de*SUMOylation limits DNR-induced-expression of the CTCF-bound *NFKB2* gene in AML.

**Figure 5:**
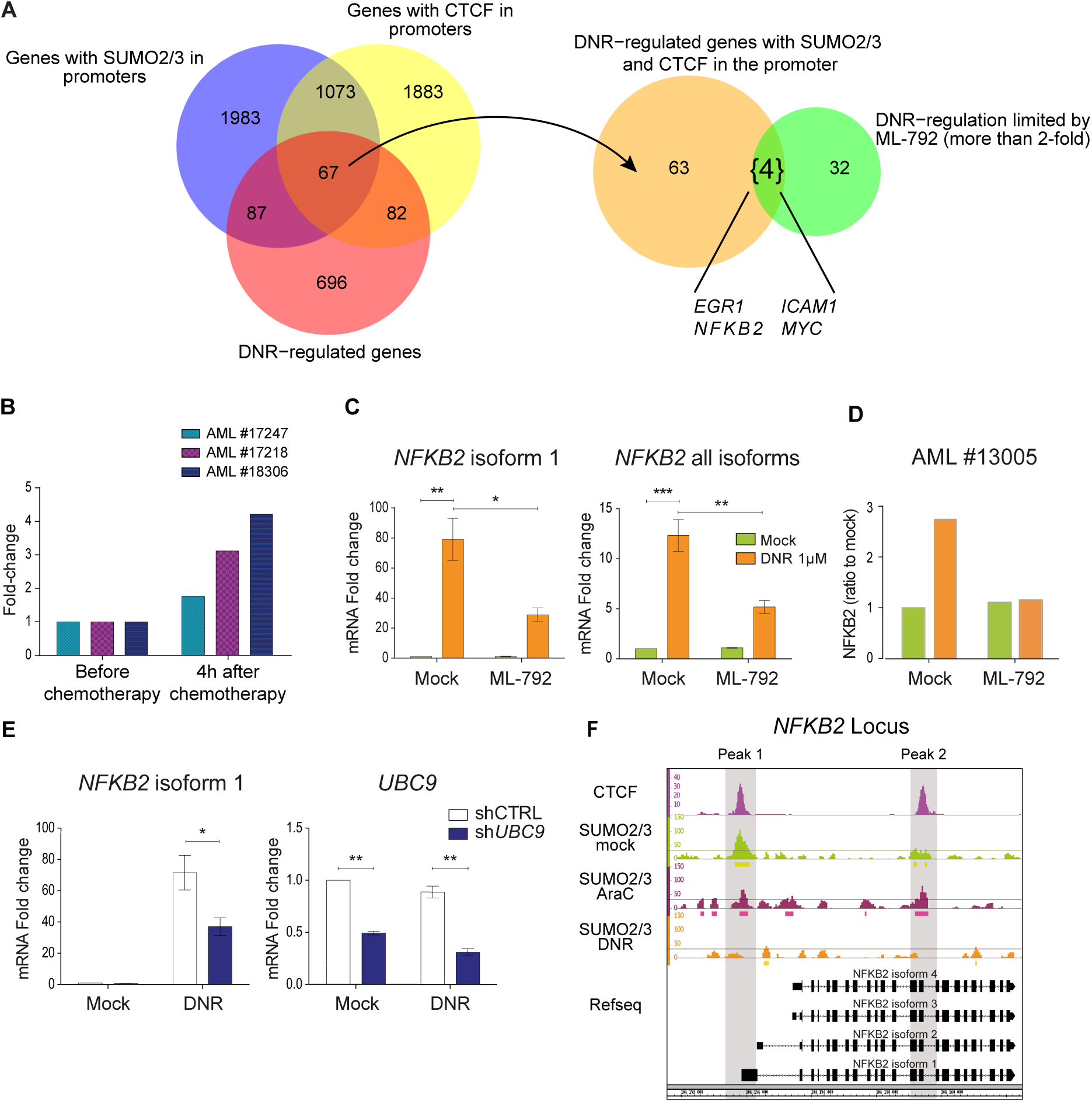
*de*SUMOylation limits DNR-induced changes in *NFKB2* expression. *A: identification of genes potentially regulated by SUMO and CTCF.* Left panel: Venn Diagram showing the intersection between the genes bound by SUMO-2/3 and CTCF in their promoter regions (−2kb-TSS) and whose expression is up- or down-regulated by DNR ≥2 fold. Right panel: Venn diagram displaying the intersection between the 67 genes bound by both SUMO and CTCF and regulated by DNR with the list of 36 genes, which their DNR-induced regulation is affected by the inhibition of SUMOylation by more than a 2-fold factor. *B: Regulation of NFκB2 gene during AML patient treatment*. Blood sample from 3 patients were collected before and 4h after the induction chemotherapy (DNR: 90mg/m^2^ and Ara-C 30mg/m^2^). PBMC were purified, mRNA prepared and *NFKB2* expression monitored by RT-qPCR, normalized to TBP and S26 levels and expressed as ratio to cells before treatment. *C, D: inhibition of SUMOylation limits NFκB2 induction by DNR.* HL-60 (C) and primary AML cells (D) were treated with 1µM of DNR for 3 hrs with or without 0.5µM of ML-792. The levels of the indicated mRNAs were measured by RT-qPCR, normalized to TBP and S26 and expressed as ratio to mock-treated cells (n=3). *E: UBC9 knock-down limits DNR-induced NFKB2 expression.* HL-60 cells stably expressing scramble or *UBC9* directed shRNA were mock- or DNR-treated for 3 hrs. The levels of the indicated mRNAs were measured by RT-qPCR, normalized to TBP and S26 and expressed as ratio to mock-treated cells (n=3). *F: CTCF and SUMO bind to the NFκB2 promoter:* ChIP-Seq data for SUMO-2/3 and CTCF were aligned and visualized using the IGB software at the level of the *NFκB2* gene.

### *De*SUMOylation limits DNR-induced chromatin 3D rearrangements at the *NFKB2* locus

Publicly available HiC data indicate that *NFKB2* is located at the center of a Topologically-Associating Domain (TAD), which extends over 500 kilobases on chromosome 10 (Figure 6A). They also suggest the existence of various long-range interactions between the *NFKB2* gene and distant regions within this TAD. Moreover, we observed that CTCF largely co-localizes with SUMO-2/3, not just at the *NFKB2* locus, but also at various places covering the whole *NFKB2* TAD in HL-60 cells (Figure 6B). Together, these observations suggested that DNR-induced *NFKB2* expression could be associated with changes in chromatin organization that could be regulated by SUMOylation/*de*SUMOylation events.

**Figure 6:**
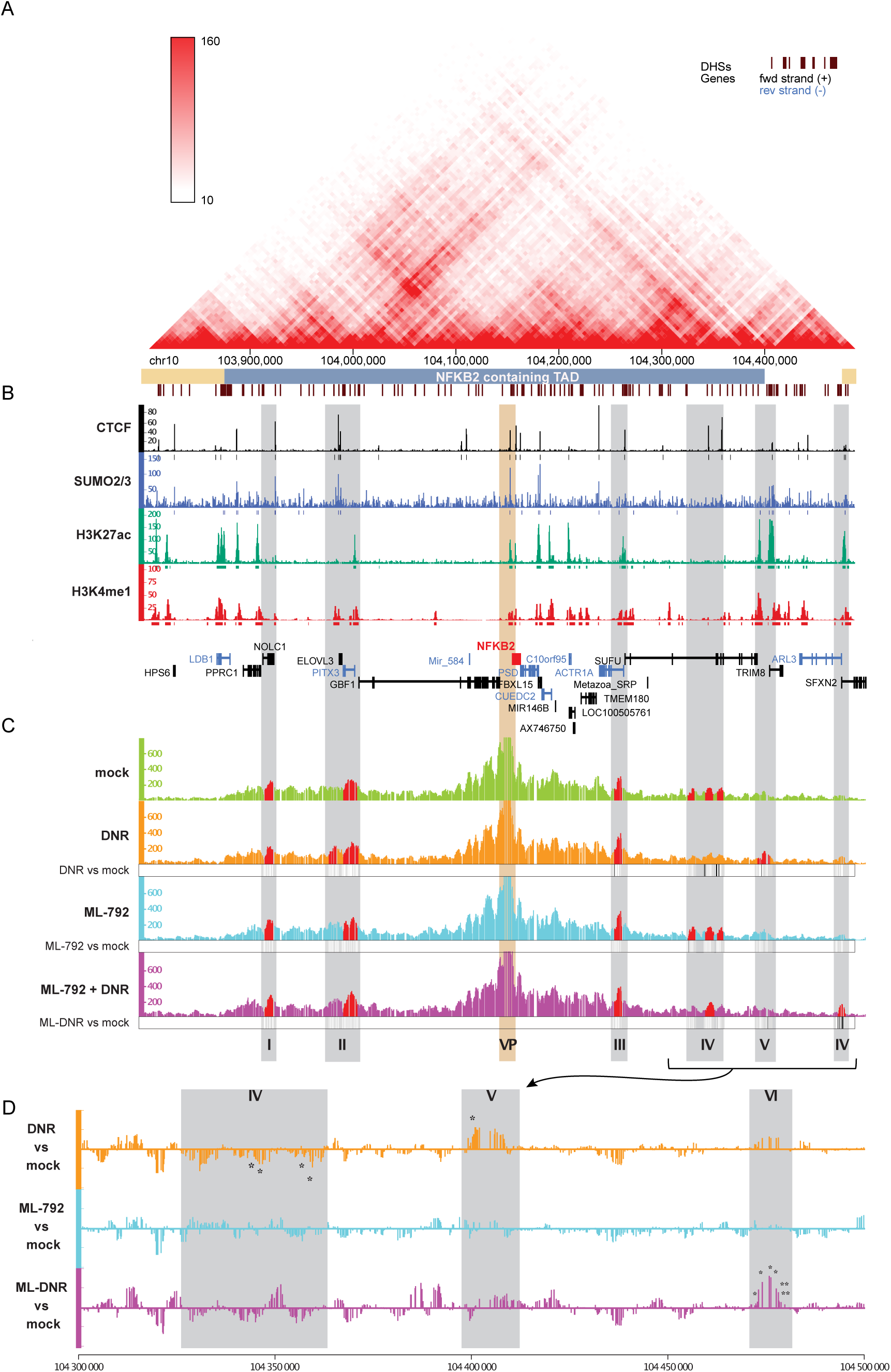
*de*SUMOylation limits DNR-induced changes in the 3D conformation of the *NFκB2* locus. *A: HiC map of the TAD containing NFκB2 gene.* This map was obtained using publicly available HiC data obtained in the K562 human chronic myeloid cell line (77). The *NFKB2*-containing TAD is underlined in blue *B: Distribution of SUMO and CTCF in the NFκB2 containing TAD.* ChIP-Seq data for CTCF, SUMO-2/3, H3K27ac and H3K4me1 are represented by the normalized read count per 50 bp bin *C,D: Inhibition of SUMOylation limits DNR-induced changes in NF-κB2 locus 3D conformation*. HL-60 treated for 2 hrs with DNR (1 µM), ML-792 (0.5 µM) or the combination and subjected to 4C experiment (3 biological replicates). The Y axis of the 4C-seq tracks represents the normalized interaction frequencies with the viewpoint (*NFKB2* promoter, VP) per 10 bp bin. Grey zones are highly reproducible interaction region in at least one condition (regions plotted in red present a *p*-value < 0.05 in the peakC analysis of the 3 replicates) and named from I to VI. D: Differential analysis of the contact point frequency in the regions IV-VI for DNR, ML-792 and ML-792+DNR compared to mock-treated cells. *p*-values for the peaks showing statistically significant differences between the conditions are indicated.

To address this issue, we resorted to Circularized Chromatin Conformation Capture (4C) experiments in HL-60 cells, using the *NFKB2* promoter as a viewpoint. In mock treated cells, we found that this promoter interacts significatively with two regions upstream of the *NFKB2* gene (regions I and II) and two downstream of it (regions III and IV) (red domains in the upper lane of Figure 6C). Noteworthy, they were all localized within the *NFKB2* TAD in the hundred kb-range from the *NFKB2* TSS and presented at least one CTCF-bound site.

The overall topology of the *NFKB2* locus was not strongly affected by a 2 hrs treatment with DNR (compare green and orange profiles in the first two lanes of Figure 6C). However, a differential profiling analysis (Figure 6D) showed decreased interactions between the CTCF/SUMO-bound *NFKB2* promoter and region IV in DNR-treated cells. Moreover, DNR induced a new interaction with a region (termed V) localized at the extreme border of the *NFKB2* TAD (Figure 6C and 6D). Interestingly, this new interacting region is enriched for histone marks (H3K27ac and H3K4me1) characteristic of active enhancers, while the interacting region in mock-treated cells (region IV) was devoid of such marks (Figure 6B). Thus, DNR-induced up-regulation of *NFKB2* is associated with changes in the frequencies of chromatin looping between its promoter region and distal regions within the *NFKB2* TAD, which include potential enhancers.

To assess whether the SUMO pathway could be involved in chromatin 3D organization changes induced by DNR, we also conducted 4C experiments on cells treated with ML-792 alone or in combination with DNR. Treatment with ML-792 alone, which did not affect *NFKB2* gene expression, did not modify the overall 4C profile of the locus (compare green and blue profiles in lanes 1 and 3 of Figure 6C and see differential profiling in Figure 6D). However, when used together with DNR, ML-792 prevented the changes observed in the presence of DNR only (*i.e.*, reduction of interactions with regions IV and induction of interaction with region V) and led to a new interaction with a region (termed VI) surprisingly localized beyond the *NFKB2* TAD border (Figure 6C and 6D). Taken together, our data suggest that *de*SUMOylation of proteins bound at CTCF-bound sites limits *NFKB2* activation by DNR by affecting the chromatin 3D architecture changes induced by DNR.

## Discussion

In this work, we report that an early effect of DNR, one of the two frontline chemotherapeutics used in AML treatment, is SUMO-controlled alteration of specific transcriptional programs. DNR modifies the expression of almost 1000 genes in chemosensitive HL-60 cells after only 3 hrs of treatment. In contrast, much less genes are regulated by Ara-C. The difference between the two drugs is not due to a lower efficacy of Ara-C to induce cell death at the dose used in our transcriptomic study. Rather, it is due to an intrinsically more limited ability of Ara-C at altering gene expression. Importantly, selected DNR-up-regulated genes were also rapidly induced by DNR in three primary AML patient samples and one of them (*NFKB2*) was also rapidly upregulated *in vivo* during standard AML chemotherapy. However, besides this, the top DNR-upregulated genes found in HL-60 cells were more induced by Ara-C than by DNR in one of the AML primary samples whereas they were hardly induced by Ara-C in the two other samples. Thus, altogether, our data indicate that DNR and Ara-C induce rapid (hour-range) transcriptome changes in AML with the effect of DNR being much stronger than those of Ara-C. However, at the same time, they also suggest a certain degree of variability that is likely explained by AML heterogeneity.

One of the main pathways we found associated with DNR-up-regulated genes is apoptosis. This suggests that the rapid gene expression changes induced by this drug set up a favorable pro-apoptotic ground that add to the DNA damages it generates for killing chemotherapy-treated AML cells at a later stage. It should, however, be noted that, in addition to pro-apoptotic genes, anti-apoptotic ones were also activated. This observation is consistent with those by others that DNR also activates pro-survival pathways such as the PI3-K/AKT- and NF-κB pathways and their targeting is considered as a potential therapeutic strategies to improve their efficiency (6, 62). Another functional category found enriched in DNR-induced genes was inflammation and immunity-related processes. In various immunocompetent mouse models, antracyclines were described as capable of inducing the immunogenic cell death of diverse solid tumors, in particular through the induction of an interferon response (64–66). The genes we identified as up-regulated in DNR-treated AML cells could participate in the development of an adaptative immune response against leukemic cells in chemotherapy-treated patients. Finally, downregulated genes are highly enriched for histone genes. This could result in decreased histone levels, which might loosen chromatin and favor the genotoxic action of the chemotherapeutic drugs. Altogether, our data suggest that the fast transcriptome changes induced by DNR before treated cells start dying may contribute to the response of AML to this drug.

Our ChIP-seq experiments showed that SUMOylated proteins are highly enriched at active promoters and enhancers in HL-60 cells, in line with data obtained in other cell types (21, 24–26). Together with our SUMOylome-wide proteomic study, they also indicated that DNR induces a massive *de*SUMOylation of chromatin-bound proteins at these places prior to DNR-induced transcriptional changes. As this occurs before massive *de*SUMOylation of cellular proteins becomes detectable (13), this indicates that DNR-induced protein *de*SUMOylation is not random in the cell. It suggests it is kinetically and spatially ordered by mechanisms that remain to be characterized (also see below). Moreover, as Ara-C, which showed much lesser effects on transcription, induces almost no change in the SUMOylation of promoter- and enhancer-bound proteins, our data suggest that DNR-induced *de*SUMOylation at these regulatory elements regulates DNR-regulated gene expression.

To address the role of *de*SUMOylation in DNR-induced gene expression alterations, we performed RNA-Seq in cells treated with DNR and/or the SUMOylation inhibitor ML-792. As DNR induces fast chromatin protein *de*SUMOylation, we first asked whether inhibition of SUMOylation alone could reproduce its effect on gene expression. This was clearly not the case as ML-792 had very small effects on gene expression (only 18 genes up-regulated and 3 down-regulated after 3 hrs or treatment). This suggests that the inhibition of SUMOylation induced by DNR is not, on its own, responsible for the fast and broad transcriptome changes. This observation is consistent with the initial report on ML-792 showing that only a very few genes are activated in cultured cells by this inhibitor, even after longer treatments (56). We therefore wondered whether protein *de*SUMOylation would have a role in the regulation of gene expression only in the presence of DNR. To this aim, we used ML-792 in combination with DNR, to accelerate and amplify the *de*SUMOylation induced by this drug. Although ML-792 had little effect on the nature and the number of the genes up- or down-regulated by DNR, it limited their up- or down-regulation, suggesting that DNR-induced protein *de*SUMOylation counteracts DNR ability to activate or repress gene expression. In particular, we found that SUMOylation inhibition reduces the activation of genes from the pro-survival NF-κB pathway, which might facilitate DNR-induced AML cell death.

Our proteomic-based study of the HL-60 cell SUMOylome characterized the proteins that are *de*SUMOylated in response to DNR. Out of the 900 SUMOylated proteins identified in mock-treated cells, only 100 were significantly *de*SUMOylated after 2 hrs of DNR treatment. Consistent with the massive loss of SUMO-2/3 observed by ChIP-seq at promoters and enhancers at the same time point, most of these *de*SUMOylated proteins are transcription factors and co-regulators. This suggests that early DNR-induced *de*SUMOylation is spatially regulated and preferentially concerns proteins bound to specific chromatin regions, many of them probably being engaged in the same protein complexes. SUMOylation stabilizes transcriptional complexes at promoters under conditions of proteotoxic stress (67), following a process called “group SUMOylation” (68). According to this concept, SUMO can control the activity of protein complexes regardless of the modified protein, or the precise sites that are SUMOylated on these proteins. It is therefore possible that DNR-induced *de*SUMOylation loosens interactions within transcription-regulating complexes binding at the promoters and/or enhancers of the genes affected by DNR, leading to changes in final transcriptional outputs. If fast DNR-induced *de*SUMOylation at precise chromatin sites is most probably partly explained by local inhibition of chromatin-bound E1 and E2 SUMOylation enzymes, it might also involve faster deconjugation of SUMO by *de*SUMOylases at these same places. For example, SENP6 was reported to *de*SUMOylate CTCF (69), one of the proteins we found *de*SUMOylated by the DNR treatment. CTCF is a multifunctional protein involved in both the regulation of chromatin 3D architecture and the control of gene expression (70). It interacts with the cohesin complex (composed of SMC1, SMC3, RAD21 and SA1/2 proteins) and is involved in the formation of diverse chromatin regulatory loops (57). Depending on the situation, such loops can activate transcription by bringing enhancers and promoters in close proximity or repress it by limiting the access of transcriptional machineries and/or regulators to gene promoters (70). CTCF is SUMOylated (58, 60), its SUMOylation being decreased by various stresses including hypoxia and oxidative stress (59). Further links between SUMO and CTCF were described on chromatin. First, the CTCF-binding consensus sequences was found enriched at genomic loci bound by SUMOylated proteins, in particular at promoters of inactive genes (25). Second, heat shock was shown to induce a transient depletion of SUMOylated proteins from CTCF-bound sites in intergenic regions and their relocation at promoters of transcribed genes (24). Third, SUMOylated proteins were found enriched at CTCF-bound sites in Drosophila and associated to enhancer blocking (71). Along this line, we found that the CTCF-binding site is the most enriched motif in SUMO-2/3 bound chromatin regions in AML cells and the co-binding of CTCF and SUMO is highly enriched at promoter and enhancers compared to intergenic regions. Moreover, the identification of CTCF as one of the proteins rapidly *de*SUMOylated upon DNR treatment, suggest that DNR-induced hypoSUMOylation of CTCF and (most probably) of other still-to-be-identified proteins present at CTCF-bound sites is involved in gene expression changes through chromatin looping alteration.

To explore this hypothesis, we focused on the *NFKB2* gene for several reasons: (i) it is one of the top-DNR-induced gene in HL-60 cells, (ii) its induction by DNR is reduced in the presence of ML-792 in, not only HL-60 cells, but also primary AML samples and (iii) its promoter region is both bound by CTCF and marked by SUMO. Moreover, it plays important roles in the control of both cell survival and inflammation/immunity (62), two of the main gene categories rapidly affected by the DNR treatment. Our 4C experiments revealed that its promoter preferentially contacts four distal regions located up to 200 kb upstream (2 regions) and downstream (2 regions) of the *NFKB2* gene, all within the *NFKB2* containing TAD and bound by CTCF in HL-60 cells. Although DNR did not markedly alter the overall architecture of the *NFKB2* locus, it induced the loss of an interaction between *NFKB2* promoter and a region devoid of active histone marks (region IV) and the appearance of a new interaction with a candidate enhancer (region V). This probably reflects the loss of a transcription-repressive loop and acquisition of a transcription-stimulating one. Consistently with its limited effects on gene expression, the sole inhibition of SUMOylation by the ML-792 inhibitor alone did not affect the overall structure of the *NFKB2* locus. This indicated that SUMOylation *per se* is not required for maintenance of the chromatin loops forming between the *NFKB2* promoter and the above-mentioned interacting regions (at least for the duration of the experiment). However, when used with DNR to accelerate/amplify DNR-induced *de*SUMOylation, ML-792 prevented the DNR-induced interaction between *NFKB2* promoter and the candidate enhancer located in region V. Instead, it induced the creation of a new interaction with a region located beyond the TAD border (region VI). This switch might prevent full activation of *NFKB2* gene. Altogether, this suggests that DNR-induced *de*SUMOylation can attenuate the transcriptional effects of DNR by controlling chromatin 3D structure. Rapid and massive changes in the SUMO proteome associated to transcriptome alterations have already been observed in response to various external cues, including heat shock (24, 67), oxidative stress (29, 72) and genotoxics such as MMS (73). Our herein data suggest that such SUMO-dependent switches might control transcriptome changes, at least in part, by affecting chromatin 3D architecture/dynamics. This is all the more to be considered that inducible genes have been reported to be more enriched in CTCF-controlled chromatin loops than housekeeping ones (74, 75). Future work will therefore have to elucidate how SUMO serves as a platform, especially at CTCF-bound sites, to recruit proteins involved in chromatin remodeling and/or structuration and how SUMOylation/*de*SUMOylation cycles at these places explain the transcriptional changes linked to alteration of 3D chromatin organization.

## Acknowledgments

We thank the members of the “Ubiquitin Family in Hematological Malignancies” group at the IGMM for fruitful discussions. Funding was provided by the CNRS, Ligue Nationale contre le Cancer (Equipe Labellisée to MP and PhD fellowships to MB), Fondation de France (2011-00025575), FRM (contract FDT20140930973 to MR and FDM201906008566 to LG). The HEMODIAG_2020 collection of clinical data and patient samples was funded by the Montpellier University Hospital, the Montpellier SIRIC and the Languedoc-Roussillon Region. MGX acknowledges financial support from the France Génomique National infrastructure, funded as part of “Investissements d’Avenir” program managed by the Agence Nationale pour la Recherche (contract ANR-10-INBS-09). Work at Novo Nordisk Foundation Center for Protein Research (CPR) is funded in part by a generous donation from the Novo Nordisk Foundation (Grant number NNF14CC0001) and the Danish Cancer Society KBVU project grant (R90-A5844).

## Notes

### Competing Interest Statement

The authors have declared no competing interest.

