## Supplementary Figures for "SUMOylation controls the rapid transcriptional reprogramming induced by anthracyclines in Acute Myeloid Leukemias"

Supplementary Figure 1

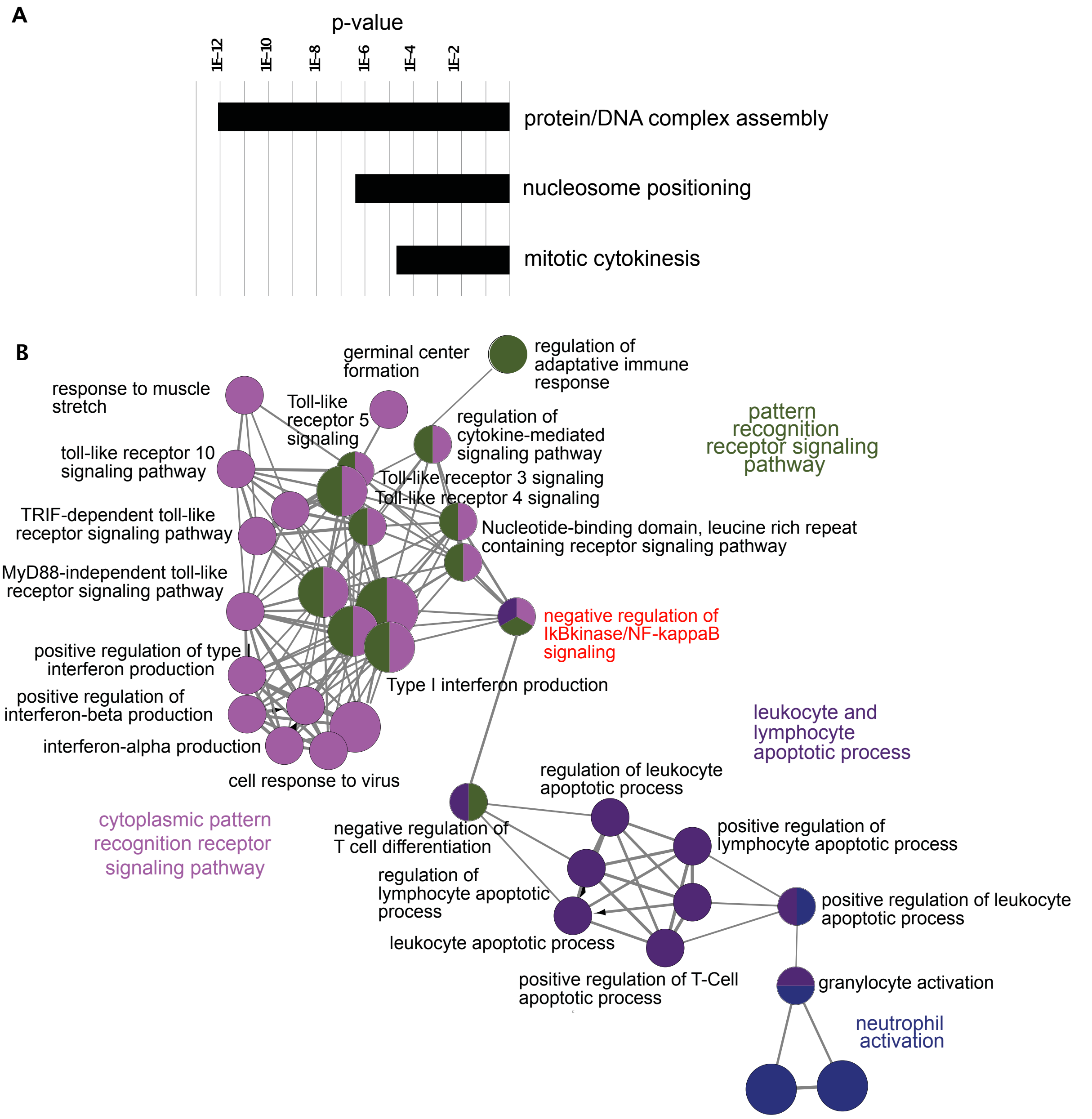


**Supplementary Figure 1: Ontology analysis of the genes regulated by DNR and Ara-C in HL-60 cells.** *A: Ontology analysis of the genes down-regulated (<-2 fold, FDR<0.05) by DNR and Ara-C.* Ontologies were obtained using the ClueGo plugin in the Cytoscape application. *B: String network of genes up-regulated by DNR*. The network was obtained with the ClueGo plugin in the Cytoscape application using all genes up-regulated (>2 fold, FDR<0.05) by DNR.

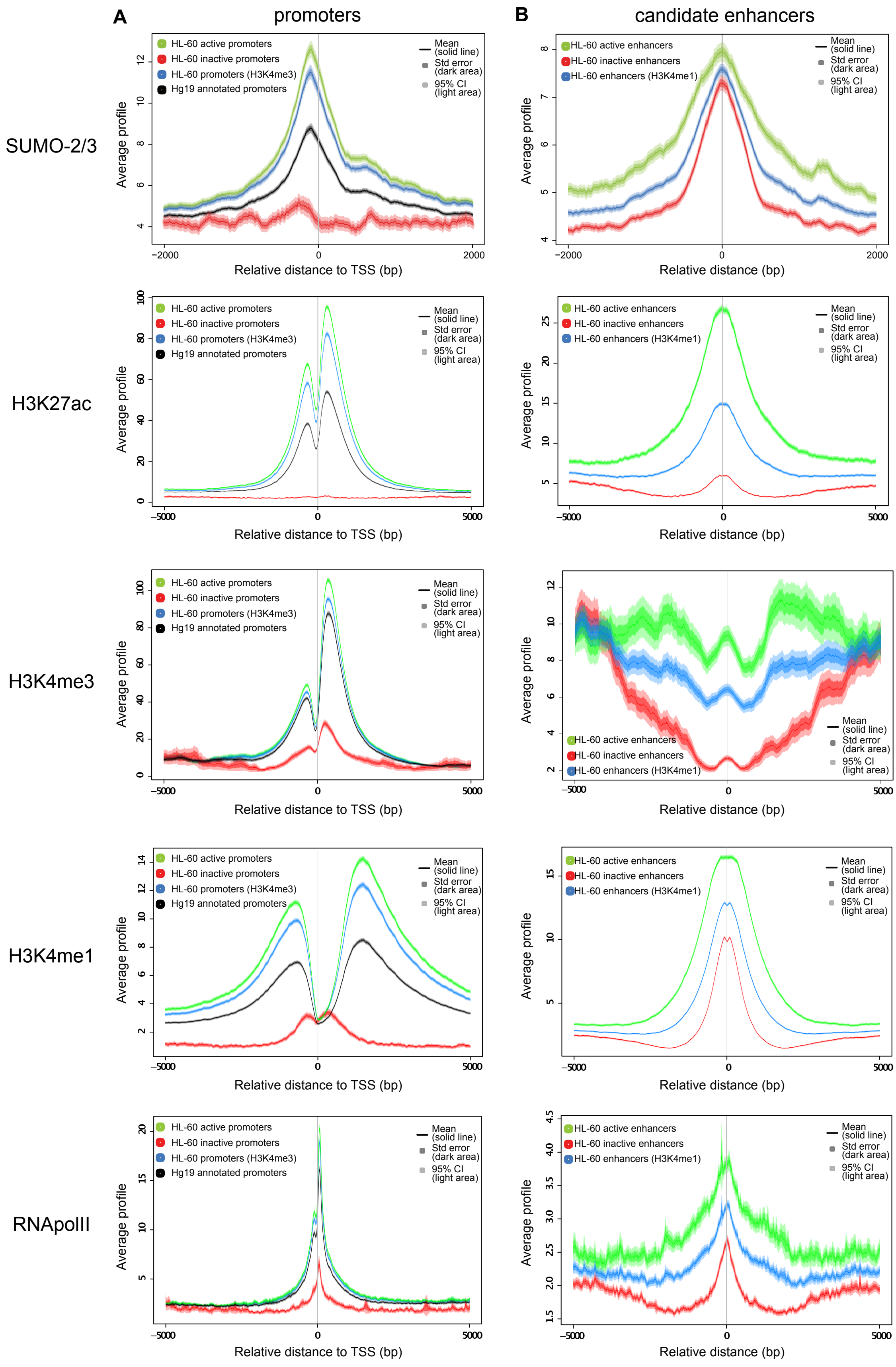


**Supplementary Figure 2: SUMO-2/3 conjugated proteins are enriched on enhancers and active promoters.** *A, B: Metaprofile of the SUMO-2/3* ChIP-seq signal on promoter (A) or enhancer (B) depending of the characteristics of *cis*-regulatory element. The promoters and enhancers as well as their level of activity were defined using publicly available histone marks ChIP-seq data (H3K27ac, H3K4me3 and H3K4me1) and RNApolII for HL-60 cells and NCBI refseq data.

Supplementary Figure 3

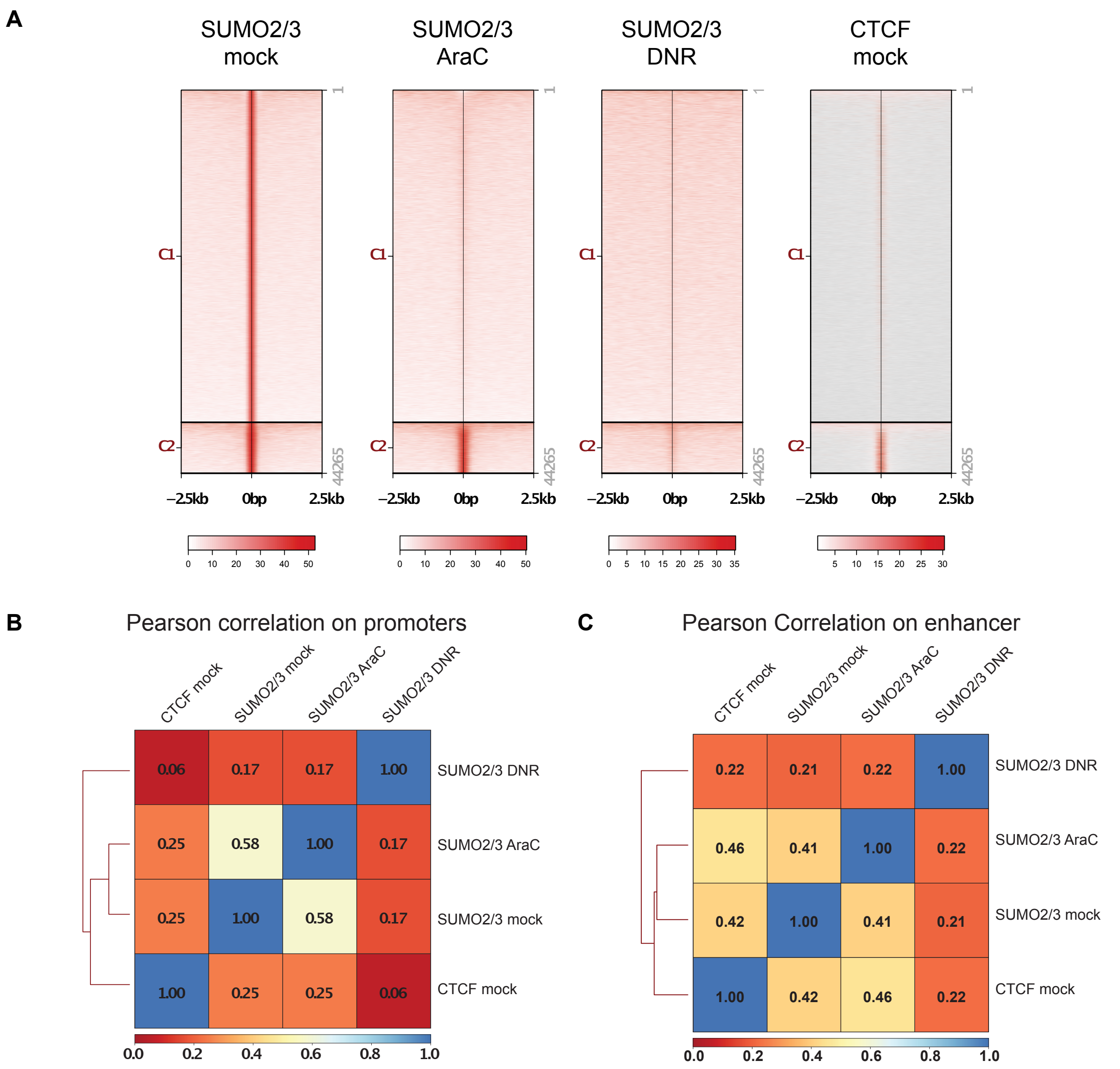


**Supplementary Figure 3: co-binding of CTCF and SUMO-2/3 on chromatin.** *A: Distribution of SUMO-2/3* ChIP-Seq peaks in Ara-C and DNR treated HL-60 cells and CTCF ChIP-Seq peaks compared to the peak region (-2.5kb; +2.5kb) of SUMO-2/3 in mock treated HL-60. *B,C: Pearson correlation matrix between ChIP-seq signals* of SUMO-2/3 in mock, Ara-C and DNR treated conditions and CTCF on HL-60 promoters (B) or HL-60 enhancers (C). Matrix were calculated with deeptool 3.1.3 (multiBigwigSummary following by plotCorrelation tools).

**Supplementary Table 1: Transcriptomic analysis of DNR and Ara-C regulated genes.**

**Supplementary Table 2: RNA-Seq analysis of the genes regulated by DNR +/- ML-792**

**Supplementary Table 3: SILAC mass spectrometry identification of SUMOylated proteins.**
